# Dynamic regulation of cystathionine gamma-lyase expression during normoxia, ischemia, and reperfusion in liver: interplay of transcription factors, microRNAs, and histone modifications

**DOI:** 10.64898/2026.10.15.759434

**Authors:** Anand Kumar Patel, Navasuja Ganesh, Vatsal Arya, Piyushkumar R. Kapopara, Gopikrishna Rathinavel, Vikas Arige, Nitish R. Mahapatra

**Author notes:** Correspondence to: Dr. Nitish R. Mahapatra, Department of Biotechnology, Bhupat and Jyoti Mehta School of Biosciences, Indian Institute of Technology Madras, Chennai 600036, India.

## Abstract

Cystathionine gamma-lyase (CTH), a key enzyme in hydrogen sulfide (H_2_S) biosynthesis, has been studied extensively in models of oxidative stress. Dysregulated CTH has been reported in cardiovascular disease, ischemia-reperfusion (I/R) injury, aging, and xenobiotic stress, where reactive oxygen species are central to pathology. Despite this, how CTH is regulated at the molecular level under basal, ischemic, and reperfusion conditions is largely unknown. We observed that CTH expression is dysregulated during ischemia and reperfusion in Huh7 hepatic cells, mouse liver, and human liver, and therefore systematically dissected its transcriptional, post-transcriptional, and epigenetic control. Promoter-deletion reporter assays identified the -275/+91 bp region as the core human CTH promoter. This region was predicted to harbour binding sites for Sp1, CREB, and NFY, which were confirmed through overexpression, knockdown or inhibition, and ChIP assays in Huh7 cells. Computational and experimental screening identified miR-150-5p and miR-510-5p as novel post-transcriptional regulators that bind directly to the CTH 3’-UTR. We also examined histone marks (H3K9ac, H3K9me3, H3K4me3, and H3K27me3) at the CTH promoter and found that H3K9ac, H3K4me3, and H3K27me3 correlate with CTH expression across basal, ischemic, and reperfusion conditions. Mechanistically, ischemia reduces Sp1 occupancy at the CTH promoter, raises miR-150-5p and miR-510-5p levels, lowers H3K9ac, collectively suppressing CTH. Reperfusion reverses this with greater Sp1 and CREB binding, lower miR-150-5p and miR-510-5p levels, and reduced H3K27me3, restoring CTH expression. Together, these findings reveal novel roles for Sp1, CREB, NFY, miR-150-5p, miR-510-5p, and histone modifications in the regulation of CTH during basal, ischemia and reperfusion conditions.

## INTRODUCTION

Cystathionine gamma-lyase (CTH) and Cystathionine beta-synthase (CBS) are the principal enzymes of the trans-sulfuration pathway and play crucial roles in sulfur amino acid metabolism and cellular redox homeostasis. CBS catalyzes the condensation of homocysteine to form cystathionine, which is subsequently converted by CTH into cysteine and hydrogen sulfide (H_2_S). In addition to maintaining homocysteine balance, both enzymes contribute to the endogenous production of H_2_S, a gaseous signaling molecule with diverse physiological functions [1]. By regulating homocysteine metabolism, cysteine availability, glutathione biosynthesis, and H_2_S production, the transsulfuration pathway serves as a critical determinant of cellular antioxidant defense and inflammatory homeostasis [2, 3]. Consequently, dysregulation of CBS and CTH has been implicated in several pathological conditions, including cardiovascular diseases, oxidative stress, inflammation, and aging [4–7].

Between these two enzymes, CTH has been widely investigated particularly because of its well-established roles in tissue protection in various *in vivo* and *in vitro* models encompassing oxidative stress as one of the underlying pathophysiology. For instance, reduced CTH expression has been reported by our group and others in hypertension [8, 9], myocardial infarction, aging-associated tissue dysfunction [5], and ischemia-reperfusion injury (IRI) [10, 11]. The critical role of CTH in cellular antioxidant defense is exemplified by the fact that genetic ablation of CTH in rodents renders these animals highly dependent on dietary cysteine supplementation for protection against damage induced by oxidative stress [12]. Moreover, the beneficial effects of calorie restriction against hepatic IR injury are linked to CTH/H_2_S signaling [13].

Following CTH loss, H_2_S supplementation has been strategically used to resolve the tissue/cellular damage caused across different oxidative stress models. In renal IR, compensating the H_2_S bioavailability was sufficient to attenuate renal damage triggered by CTH deficiency [14]. Similarly, H_2_S administration protected kidneys from hypoxia-induced damage by inducing a hypometabolic state [15]. In addition, xenobiotic compounds such as diethylnitrosamine (DEN) and carbon tetrachloride (CCl4), induce excessive free radical formation in liver cells are widely employed in oxidative stress models [16–20]. Notably, CTH-deficient mice exhibit heightened susceptibility to DEN- and CCl4-induced hepatic injury, characterized by increased lipid accumulation, dysregulated lipogenesis, and exaggerated inflammatory responses. Importantly, exogenous H_2_S supplementation effectively alleviates liver damage induced by these agents [21–24].

These observations not only highlight the indispensable role of the CTH/H_2_S axis in maintaining tissue resilience under oxidative stress conditions but also suggest that H_2_S supplementation may serve as a therapeutic strategy to tackle oxidative stress related injuries. However, a major limitation of current H_2_S donors is their short half-life and rapid, poorly controlled H_2_S release, which limits their suitability for therapeutic applications. A complementary strategy is to modulate endogenous H₂S production by cystathionine γ-lyase (CTH) on the basis of clinical assessment. For instance, transgenic overexpression of CTH significantly reduced myocardial ischemic injury in mice [6]. Over the past few years, we and others have identified several molecular regulators governing CTH expression under basal and various pathological conditions [8, 25–33]. However, due to the complexity of gene regulation processes the mechanism of regulation of CTH remains incompletely understood.

We sought to define the regulatory network governing human CTH expression in hepatocytes by identifying transcription factors, miRNAs, and epigenetic mechanisms including histone modifications that control its basal expression. We then interrogated how these regulatory determinants shape CTH expression during hepatic ischemia and reperfusion, considered as distinct pathophysiological states. Our comprehensive computational and experimental analyses revealed previously unrecognized roles of three transcription factors (viz. CREB, NFY and Sp1), and two miRNAs (viz. miR-150-5p and miR-510-5p) that regulate CTH expression under basal, ischemia, and reperfusion conditions in the liver. Additionally, we have also identified epigenetic markers H3K9ac, H3K4me3 and H3K27me3 that contribute to the dynamic regulation of CTH expression under these conditions. Collectively, these findings establish a mechanistic framework for precision modulation of the CTH-H₂S axis, with translational potential to guide the development of targeted therapeutic strategies for hepatic IRI.

## MATERIALS AND METHODS

### Generation of human CTH promoter-luciferase reporter constructs, CTH-3′-UTR-luciferase reporter constructs, and microRNA expression plasmids

Varying lengths of proximal CTH promoter (viz. -816, -616, -375, -275, -123 and -75 bp to +92 bp) regions were amplified using the human genomic DNA sample from a healthy volunteer as a template, Phusion High-Fidelity DNA polymerase (Finnzymes, USA) and primers (Table S1) specific to the CTH promoter (NCBI reference number: NM_001902.6). PCR-amplified DNA segments were cloned at *Kpn*I and *Xho*I sites in the promoter-less firefly luciferase reporter vector pGL3-Basic (Promega, USA) (Fig. S1 a and b). These constructs were named hCTH-pro-816, hCTH-pro-616, hCTH-pro-375, hCTH-pro-275, hCTH-pro-123, and hCTH-pro-75. All the constructs were generated by PCR amplification, followed by restriction digestion with *Kpn*I and *Xho*I restriction enzymes, gel purification, and ligation. Similarly, the CTH 3′-UTR region was amplified using the human genomic DNA as template, Phusion High-Fidelity DNA polymerase and CTH 3′-UTR specific primers (Table S2). The resulting amplicon was cloned at *Xho*I and *Hind*III sites in the pmiR-REPORT vector (Life Technologies, USA). The hsa-miR-510-5p expression plasmid was generated by PCR amplification of pre-miRNA-510 region using human genomic DNA as a template, Phusion High-Fidelity DNA polymerase (Finnzymes, USA) and primers (Table S3). The PCR products were cloned at *BamHI* and *XhoI* sites of the pcDNA3.1 vector (Invitrogen, USA). The sequence and orientation of all these plasmid constructs were confirmed by Sanger sequencing. The plasmid DNA constructs were purified by HiPurA Endotoxin-free plasmid DNA Midiprep purification kit (Hi-Media, India) for transfection experiments.

### Generation of mutant CTH promoter- and 3′-UTR luciferase reporter constructs

Mutations were generated at the Sp1 and CREB and NFY-A binding sites by site-directed mutagenesis using wild-type (WT) human CTH promoter-reporter plasmid as template, primer pairs with the desired mutations (Table S4) and Phusion High-Fidelity DNA polymerase (Finnzymes, USA). Similarly, mutations in the miR-150-5p and miR-510-5p binding sites were made via deletion of those sites in 3′-UTR-luciferase constructs using WT human CTH 3′-UTR plasmids as template and specific primer pairs (Table S5). The PCR products were subjected to DpnI digestion to digest the template plasmids The digested products were used for bacterial transformation, followed by identification of mutant clones. The accuracy of all the mutant plasmid constructs was confirmed by Sanger sequencing. These plasmid DNAs were amplified and purified by HiPurA Endotoxin-free plasmid DNA Midiprep purification kit (Hi-Media, India) for transfection experiments.

### *In silico* analysis of human CTH core promoter and CTH 3′-UTR regions

The putative transcription factor (TF) binding sites in the CTH core promoter (viz., -275 to +1-bp region, which displayed the highest reporter activity) were identified using multiple TF prediction tools such as Alibaba [34], Consite [35], Matinspector [36], Lasagna [37], TF bind [38], and Promo [39]. Similarly, multiple miRNA-prediction tools such as mirDB [40], TargetScan [41], miRwalk [42], miRNAMap [43], microT-CDS [44], EIMMo [45] were used to predict the putative miRNAs that might interact with the CTH 3′-UTR (Table S6). These *in silico* analyses revealed putative binding sites for several TFs and miRNAs. TFs predicted by at least two programs and miRNAs predicted by at least four programs were short-listed for experimental validation.

### Cell culture and transient transfections

Kidney HEK-293, neuronal N2a, liver Huh7, cardiomyocyte H9c2 and endothelial EA.hy926 cell lines were obtained from the Indian National Repository for Cell Lines and hybridomas at the National Centre for Cell Sciences, Pune, India. Cells were cultured in Dulbecco’s modified Eagle’s medium (DMEM) with high glucose and glutamine (Gibco, USA), supplemented with 10% fetal bovine serum (Gibco, USA), penicillin G (100 U/mL), and streptomycin sulfate (100 mg/mL) (Invitrogen, USA) in T-25 tissue culture flasks (Tarsons, India) at 37 °C with 5% CO_2_. These cell lines were routinely tested for mycoplasma infection and treated with BM-Cyclin (Merck, USA) to eliminate mycoplasma, if detected. Transient transfections of these cell lines with the CTH promoter-luciferase reporter constructs (WT and mutated), 3′-UTR -luciferase reporter constructs (WT and mutant) (400 ng per well of a tissue-culture-treated 24-well plate) were carried out using polyethyleneimine (PEI, Sigma). A β-galactosidase expression plasmid (400 ng per well) was co-transfected as an internal control for normalization of transfection efficiency across different wells [46] For experiments involving overexpression/downregulation/inhibition of TFs, the CTH core promoter-reporter construct (*viz.*, CTH-pro-275; 400 ng/well) was co-transfected with increasing doses of Sp1 expression plasmid containing the human full-length Sp1 cDNA fused to the CMV promoter (obtained from Dr. Dona Lee Wong, Harvard Medical School, Boston [47]) or CREB expression plasmid (CREB-VP16, which contains basic-leucine zipper domain of CREB, and the transcriptional activation domain of the herpes virus VP16 protein (obtained from Dr. David Ginty, Howard Hughes Medical Institute) [48] or with Sp1 shRNA plasmid (obtained from Dr. Guido Marcucci, Ohio State University, Ohio) [49] or KCREB (a dominant repressor of CREB, which has a single mutation, Arg to Leu, in the DNA binding domain compared to WT (obtained from Dr. Richard H. Goodman, Oregon Health Sciences University, Portland, [50]) into Huh7 cells. As NFY is a heterotrimeric TF composed of NFY-A, NFY-B, and NFY-C subunits, a combination of three expression plasmids, each having cDNA for NFY-A, NFY-B, and NFY-C subunits were used. Dominant negative form of NFY-A (NFYDN) was used for NFY inhibition experiments. NFYDN expression plasmid expresses a dominant-negative form of NFY-A that harbors a mutation in the DNA binding domain and thus abolishes the binding of NFY trimer on CCAAT boxes on promoter. NFY-A/B/C and NFYDN expression plasmids were obtained from Dr. Shioko Kimura, National Institutes of Health, Bethesda, Maryland [51]. Total protein was used as normalization control across different wells for TF overexpression or knockdown/inhibition studies.

### Firefly luciferase reporter and ß-galactosidase assays

Transfected cells were lysed using a phosphate buffer supplemented with 0.1% Triton X-100 and 10 mM dithiothreitol on a shaker for 10 minutes. For each well, 60 μL of the clear lysate was used to record the firefly luciferase activity using an automated luminometer (LB-953, Berthhold, Germany) and luciferase buffer (1M Tris Acetate, 1M Magnesium Acetate, 0.5M EDTA, 200 mM ATP, 10 mM D-Luciferin) as described previously [52]. For β-gal assay, the cell lysate and ß-gal buffer in a ratio of 1:1 and incubated at 37^0^ C for 2 hr. The absorbance was recorded at 415 nm using a microplate UV-vis spectrophotometer [53]

### Western blotting

In order to assess the over-expression/ down-regulation/ inhibition of transcription factors and their effects on endogenous CTH levels, Huh7 cells (seeded in 24-well plates) were co-transfected with CTH promoter-reporter construct (400 ng/well) and increasing doses of CREB, Sp1 or NFY-A/B/C expression plasmids or with increasing doses of KCREB, Sp1 shRNA, or NFY-DN expression plasmid using PEI transfection reagent. Similarly, to ascertain the effect of over-expression of miRNAs on 3′-UTR activity and endogenous levels of CTH, CTH-3′-UTR reporter construct (400 ng/well) was co-transfected with increasing concentrations of hsa-miR-150-5p (pcDNA3.2/V5 hsa-mir-150 (purchased from Addgene; plasmid # 26310) or hsa-miR-510-5p expression plasmids using PEI reagent. In another set of experiments, control oligo, miR-150-5p, and miR-510-5p inhibitors (IDT, USA) were transfected into Huh7 cells using Lipofectamine 3000 transfection reagent (Invitrogen, USA). In each case, 30 h post-transfection, cells were lysed in RIPA buffer [50 mM Tris-HCl (pH 7.2), 150 mM NaCl, 1% (v/v) Triton X-100, 1% (w/v) sodium deoxycholate and 0.1% (w/v) SDS] with PMSF (Sigma, USA) and protease inhibitor cocktail (PIC) (Sigma, USA) for Western blot analysis. To isolate protein from mouse liver tissues, the tissues were minced in RIPA buffer supplemented with PIC (0.5% v/v) and PMSF (1% v/v) using a mechanical homogenizer followed by sonication. The protein concentration was estimated using the Bradford’s reagent (Hi-Media, India). SDS-PAGE was performed using a 10% SDS-polyacrylamide gel. The proteins were transferred onto nitrocellulose or PVDF membranes. A pre-stained protein ladder (Abcam, USA) was used as a molecular weight marker. Either 3% BSA or 5% skimmed milk solution prepared in Tris-buffered saline (TBS) supplemented with 0.05% Tween 20 (Hi-Media, India) (henceforth, TBST) was used for blocking the membranes for 1 hr at room temperature. Specific primary antibodies such as CTH (Cell Signaling Technology, USA, 19689) at dilution 1:2500, CREB (Cell Signaling Technology, USA, 9179) at dilution 1:2000, NFY-A (Santacruz Biotechnology, USA, sc-10779) at dilution 1: 1000, and Sp1 (Santacruz Biotechnology, USA, sc-17824) at dilution 1: 2500 were prepared in 3% BSA solution (in TBST). The blocked membranes were then incubated overnight at 4^0^C with different primary antibodies. The next day, these membranes were washed 3 times (10 minutes each) using TBST under shaking conditions. The membranes were then incubated with Horse radish peroxidase (HRP)-conjugated secondary antibody specific for either rabbit (Jackson Immunoresearch, USA, 111-035-003) or mouse (Jackson Immunoresearch, USA, 115-035-003) at 1:5000 dilution, prepared either in 3% BSA or 5% skimmed milk solution in TBST. The protein bands were detected using enhanced chemiluminescent (ECL) reagent (BioRad, USA) and iBright CL1000 imaging system (Thermofisher, USA). The intensities of bands were quantified using iBright image analysis software from Thermofisher.

### Chromatin immunoprecipitation (ChIP) assays

ChIP assays for assessment of interactions between promoter regions and TFs were carried out as described earlier [54] Briefly, Huh7 cells were fixed using 1% formaldehyde, lysed, and nuclei pellets were collected by centrifugation and re-suspended in the nuclei lysis buffer. The chromatin preparation was fragmented using a Bioruptor Pico® (Diagenode, Belgium). Sheared chromatin was used for subsequent immunoprecipitation. Pre-cleared chromatin (using protein-G-coated Sepharose beads (Invitrogen, USA, 101242) supernatants were subjected to immunoprecipitation at 4 °C overnight using 4 μg ChIP-grade antibodies against Sp1 (Santa Cruz Biotechnology, sc-17824), CREB (Cell Signaling Technology, 48H2), or NFY-A (Santa Cruz Biotechnology, USA, sc-17753) in 1x ChIP buffer supplemented with PIC. As a negative control, an equal amount of chromatin was precipitated with a pre-immune IgG antibody (Sigma-Aldrich). Specific antibody-chromatin complexes were then precipitated by incubation with ChIP-grade protein-G-coated Sepharose beads (Invitrogen, USA, 101242) for 4 hr at 4^0^C. Supernatant was collected and used for reverse cross-linking protein/DNA complexes to free DNA. The enrichment of the CTH promoter in the purified DNA was determined by real-time PCR using Kappa Sybr® Fast RT PCR kit (Roche, Switzerland, KK4602) with specific primer pairs encompassing the transcription factor binding domain (Table S7). Respective IgG enriched promoter values were used for normalization of IP data, which were then normalized with input and represented as % Input. ChIP assays for detection of histone modification marks were also performed in the similar manner. The chromatin fragments were immunoprecipitated with antibodies against H3K4me3 (Cell Signaling Technology, C42D8), H3K9me3 (Cell Signaling Technology, D4W1U), H3K27me3 (Cell Signaling Technology, C36B11), H3K9ac (Cell Signaling Technology, C5B11). Promoter enrichment was estimated by RT-PCR using multiple primer pairs (Table S8) encompassing 472 bp promoter upstream to transcription start site (TSS).

### Liver ischemia and reperfusion in an animal model

The C57BL/6 mice (male, age: 10-12 weeks old) were procured from Mass Biotech, Chennai and were housed in the animal house facility at the Indian Institute of Technology Madras. The mice were fed an ad libitum diet and were subjected to 12-hour light and dark cycles. Before surgery, the animals were subjected to overnight fasting. During surgery, the mice were injected with analgesics (meloxicam: 2mg/kg) and were anaesthetized with isoflurane (3-5% for induction and 1-2% for maintenance) and immobilized on the platform maintained at 37 ^0^C throughout the surgical procedure. The liver ischemia (∼70%) was induced by clamping the portal triad for 35 minutes using a metallic micro clamp as described previously (cite reference). The blockade in the blood supply was confirmed by checking the blanching of the liver. After the ischemia period, the clamp was carefully removed, and restoration of the blood supply was confirmed by a change in the colour of the liver. Following this, the abdomen was sutured back, and animals were brought to consciousness by flushing pure oxygen until the animal regains consciousness. After 6 hr (counted just after removing the clamp), the animals were sacrificed by over-anesthetizing using isoflurane. Blood and liver tissues were harvested and stored at -86 ^0^C. The animal experiments were approved by the Institute Animal Ethics Committee of Indian Institute of Technology Madras (Approval number: IITM/IAEC/NRM-2/05/2023).

### Ischemia-reperfusion in Huh7 cells *in vitro*

Ischemic condition was created *in vitro* by depriving the Huh7 cells of nutrients and oxygen. Briefly, the cells were seeded in a 60 mm culture dish up to maximum of 70 % confluency in DMEM supplemented with 10% FBS. During ischemia, the culture media was replaced with oxygen glucose deficient media (OGD) [124 mM Sodium Chloride, 4 mM KCl, 2 mM CaCl_2_, 1.5 mM MgSO_4_, 26 mM NaHCO_3_, 1.5 mM NaH_2_PO_4_, 10 mM Sucrose] and the dishes were kept in an airtight chamber which was purged and filled with 95% Argon and 5% CO_2_ for 2 min with airflow at 10 (unit) for 2 minutes. The cells were then incubated at 37 ^0^C. For only ischemia group, the cells were lysed immediately after they were taken out from the air-tight chamber using RIPA buffer. For reperfusion, the OGD was replaced with DMEM but without serum and cells were incubated at 37^0^C with 5% CO_2_. After reperfusion, the cells were lysed. Total protein and RNA were isolated from the cell lysates using our standardized protocols [46]

### RNA isolation from Huh7 cells and mouse liver tissue samples, and quantitative real-time PCR

Total RNA was isolated from Huh7 cells using RNAiso plus reagent (Takara, Japan, 9109). cDNA was synthesized using 2 μg RNA, random hexamers and High-Capacity cDNA Reverse Transcription Kit (Takara, Japan). Similarly, total RNA was isolated from mouse liver tissue samples using RNAiso plus reagent and NuceloSpin RNA/Protein kit (Macherey-Nagel, USA) and was subjected to cDNA synthesis. In order to determine the mRNA levels, RT-PCR was carried out using Kappa Sybr® Fast RT-PCR kit and CTH transcript-specific primer pairs (Table S9). The mRNA levels of β-actin were quantified for normalization of gene expression using actin-specific primer pairs (Table S10). In order to quantify the miRNA levels, total RNA was isolated from Huh7 cells/liver tissues as mentioned above. Two micrograms of RNA samples were used to synthesize cDNA using a high-capacity cDNA synthesis kit, miR-150/miR-510 and U6-specific stem-loop reverse transcription primers (Table S10). RT-PCR was carried out using Kappa Sybr® Fast RT PCR kit, miR-150/miR-510 and U6 forward primers, and universal reverse primer The relative abundance in each sample was determined by calculating 2^(−ΔΔCt)^ as described previously [55].

### Data presentation and statistical analysis

For CTH promoter or 3′-UTR-reporter transient transfections, co-transfection experiments were carried out at least three times, and results were expressed as mean ± SD of triplicates from representative experiments. Statistical significance was determined by Student’s *t*-test (unpaired, two-tailed) and one-way ANOVA multiple comparisons post-test, as appropriate, by using the GraphPad Prism 10 software (GraphPad Software, USA). The graphical figures were generated using BioRender.

## RESULTS

### CTH expression is dysregulated during ischemia and reperfusion

Given the central role of CTH in the transsulfuration pathway and its contribution to both cysteine metabolism and endogenous H₂S production (Fig. 1 a), we first investigated whether CTH expression is dynamically regulated in response to ischemia-reperfusion. We chose hepatic cells as those express the highest level of CTH among various cell types. In Huh7 cells, CTH mRNA levels decreased approximately by 2-fold (P = 0.0024) following 4 hr of ischemia as compared to normoxic controls (P = 0.0024) (Fig. 1 b). In contrast, after 1 hr of reperfusion, CTH mRNA levels increased approximately by 4-fold (P<0.0001) relative to ischemia (Fig. 1 b). Consistent with the changes observed at the transcript level during ischemia, endogenous CTH protein levels decreased approximately by 2-fold following 4 hr of ischemia (P = 0.0032) (Fig. 1 c). However, CTH protein levels did not recover after 1 hr of reperfusion (P = 0.5544) (Fig. 1 c), suggesting a delay between the transcriptional response and changes in protein abundance. To examine the kinetics of this response, we extended the reperfusion period to 6 hr and observed a time-dependent increase in CTH protein levels during reperfusion (Fig. 1d).

**Fig. 1.**
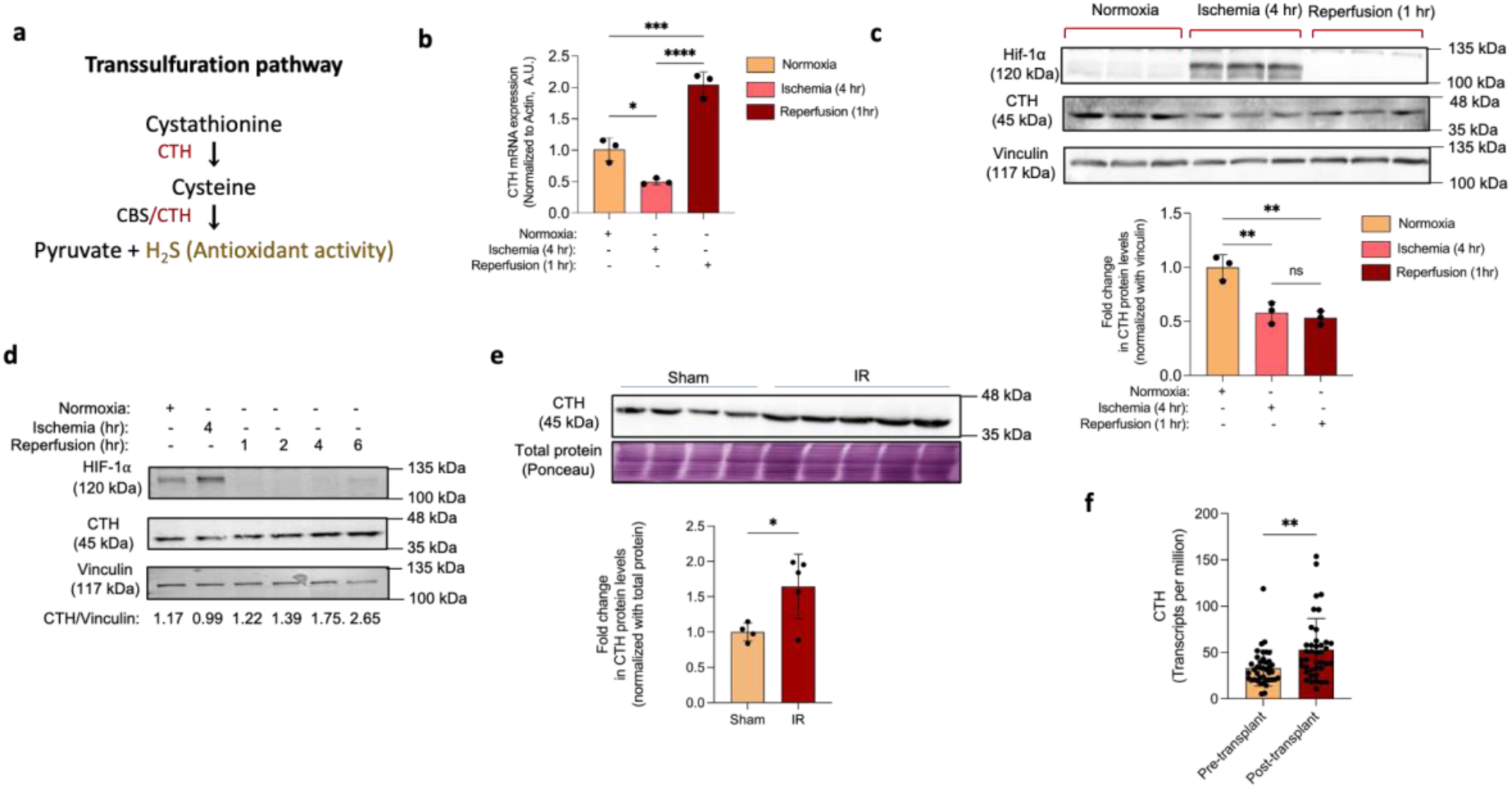
Differential expression of CTH during normoxia, ischemia and reperfusion. (a) Transsulfuration Pathway: Cystathionine (generated from homocysteine by cystathionine beta synthase [CBS]) is converted to cysteine by CTH; CTH/CBS convert cysteine to H_2_S. (b) Endogenous CTH mRNA levels in Huh7 cell line normoxia, ischemia (4 hr) and reperfusion (1 hr). (c) Confirmation of ischemia by Hif-1*a* protein presence, and endogenous CTH protein levels in Huh 7 cells challenged with IR. (d) Changes in the endogenous CTH protein levels with increasing reperfusion time (1 hr, 2 hr, 4 hr, and 6 hr) after 4 hr of ischemic insult. (e) CTH protein levels in mouse liver tissue in sham (control, n = 4) and IR group (n = 5) (f) CTH transcripts levels (transcripts per million) from human liver tissues used for orthotopic liver transplant. Pre-transplant (n=40) and post-transplant (n=40) CTH transcript values correspond to the same tissues. The values in the graph are mean ± S.D of n = 3 biological replicates. The statistical significance was determined by unpaired *t*-test or one-way ANOVA with Tukey’s multiple correction post-hoc test. ****P<0.0001, **P<0.01, *P<0.05, ***P<0.001, ns: not significant

To corroborate these *in vitro* findings, we next investigated whether this response was also observed *in vivo* using a mouse model of hepatic ischemia-reperfusion injury. Indeed, following 35 min of warm ischemia and 6 hr of reperfusion, hepatic CTH protein levels were ∼1.8-fold higher than those in normoxic control livers (P = 0.0317) (Fig. 1 e). Moreover, to extend the relevance of these findings in human liver, we analyzed a publicly available RNA-sequencing dataset from the Gene Expression Omnibus (GEO) (accession ID: GSE151648 (Table S11). CTH expression was ∼1.5-fold higher in human liver tissue after 6 h of reperfusion as compared to healthy liver tissue (P = 0.0018) (Fig. 1 f). Overall, these findings demonstrate dynamic regulation of CTH during hepatic ischemia-reperfusion, characterized by an initial reduction during ischemia followed by increased expression during reperfusion. This response was observed across cultured hepatic cells, murine liver, and human liver transcriptomic database, supporting the association of CTH regulation with the hepatic ischemia-reperfusion response.

### Identification of the core-promoter domain of the human CTH gene

Although the promoter regions may span up to several hundreds of bases upstream to the transcription start site, studies show that a core promoter is usually associated with recruitment of transcriptional machinery and basal transcription [56–58]. Therefore, we sought to identify the core promoter domain of the human CTH gene and TFs that could govern CTH expression under basal, ischemia, and reperfusion conditions.

Analysis of the 1000 bp human CTH proximal promoter (sequence retrieved from the UCSC genome browser) was carried out to identify binding sites for different TFs using multiple *in silico* tools. Based on the locations of putative TF binding sites, varying lengths of CTH promoter segments (viz., -816 bp, -616 bp, -375 bp, -275 bp, -123 bp, -75 bp to +91 bp) were PCR-amplified in such a way that the binding sites of one or more TFs were deleted between successive deletion constructs (Fig. 2a). Thereafter, these deleted promoters were cloned upstream of the promoter-less pGL3-Basic luciferase reporter vector (Fig. S1 a and b), and the basal activities of the promoter-reporter constructs were studied in Huh7, HEK-293, EA.hy926, H9c2, and N2a cells. The CTH -275 bp/+91 bp promoter construct*, in general,* displayed significantly higher/similar activity as compared to other constructs-across the cell lines. (Fig. 2 b-f)). A further deletion of the CTH promoter to -75 bp decreased the promoter activity by ∼1.5 to 2-fold in all the tested cell lines. These observations suggest that the promoter region between -275 and -75 bp potentially harbour binding sites for transcriptional activators that may be crucial for CTH expression in various cell types. Therefore, the hCTH-pro-275 promoter-reporter construct (core promoter region) was used for all the subsequent investigations.

**Fig. 2.**
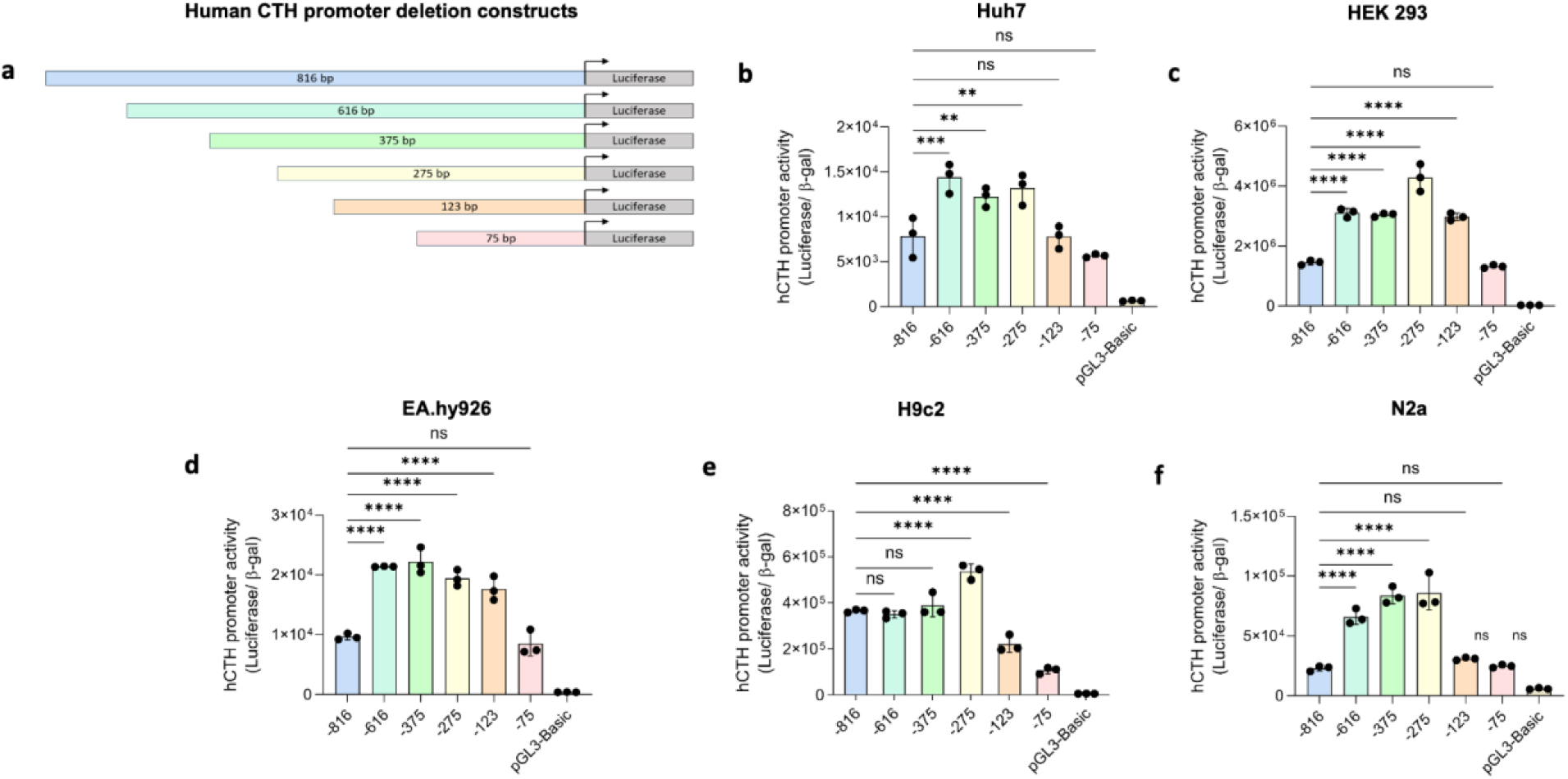
Activities of the human CTH promoter -reporter constructs in various cell lines. (a) Schematic representation of varying lengths of CTH promoter cloned into pGL3-Basic vector. The deleted promoters were cloned upstream to firefly luciferase reporter gene. (b-f) Human CTH promoter-reporter activity was recorded after co-transfection of promoter-reporter deletion constructs with a pCMV-β-galactosidase expression plasmid as transfection control in Huh7, HEK 293, EA.hy926, H9c2, and N2a cell lines. The luciferase reporter activity was normalized with β-galactosidase activity. The values in the graphs are mean ± S.D of n = 3 technical replicates. The transfection experiments were repeated at least three times and representative data are shown. The statistical significance was determined by one-way ANOVA with Tukey’s multiple correction post-hoc test. ****P<0.0001, ***P<0.001, **P<0.01; ns: not significant

### Putative TF binding sites in the human CTH promoter and correlation analysis between the transcript levels of the predicted TF and CTH across human tissues

A stringent binding motif analysis using TF prediction tools (four or more) consistently predicted binding sites for Sp1, CREB, AP-2α and NFY-A in the human CTH core promoter region (Fig. 3 a and b). The binding scores mentioned for the four TFs (NFY-A: 13.7; CREB: 11.438; AP-2α: 9.078; Sp1: 6.624) (Fig. S2 a-d) are from the ConSite TF prediction tool. Next, to investigate the likelihood of these TFs in driving/governing CTH expression in human tissues, we used existing publicly available tissue expression data and performed correlation analysis between the expression of individual TFs and CTH. Using expression data from the GTEx Portal (Table S12), a significant positive correlation was observed between CREB and CTH (r = 0.4927, P = 0.0003), Sp1 and CTH (r = 0.3066, P = 0.0322), and NFY-A and CTH (r = 0.6123, P<0.0001) across diverse human tissues; however, AP-2α, despite exhibiting a higher predicted binding score, showed no significant correlation with CTH expression (r=-0.2809, P = 0.0506) (Fig. 3 c-f). Furthermore, analysis of liver transcriptomics data from the GEO Dataset (accession:GSE151648) (Table S13) revealed a similar correlation between these TFs and CTH (AP-2α: r = -0.1943, P=0.2360; CREB: r = 0.4167, P = 0.0083; Sp1: r = 0.5323, P = 0.0005; NFY-A: r = 0.4913, P = 0.0015) (Fig. 3 g-j). In addition, AP-2α/CREB/NFY-A/Sp1 binding motif analysis using mouse CTH promoter revealed multiple binding sites for CREB, Sp1 and NFY-A but not for AP-2α (Fig. S2 e). Hence, AP-2α was excluded from further investigations. Taken together, these observations provide compelling evidence for crucial role of CREB, Sp1 and NFY in CTH expression across species.

**Fig. 3.**
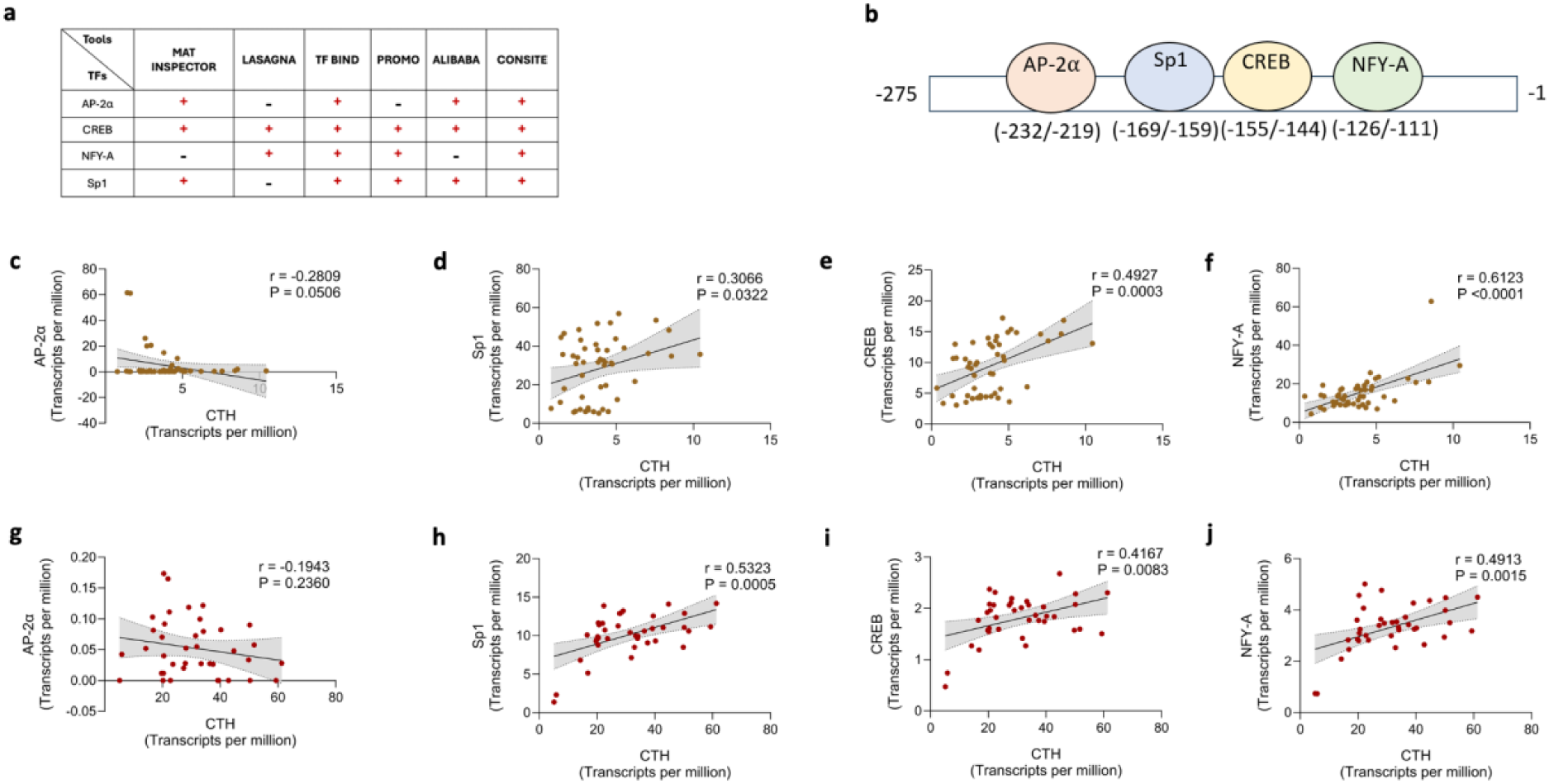
Prediction of putative TFs for regulation of the human CTH gene. (a) Six TF prediction tools were used for prediction of TFs that might bind to the core promoter region of human CTH. TFs that were predicted by four or more tools were selected for further analysis. (b) Schematic representing the binding sites of AP-2α, Sp1, CREB and NFY-A on the core promoter of human CTH gene. (c-f) The co-expression correlation analysis between the CTH and TF transcripts in healthy human tissues using normalized expression data (transcript per million) for TFs and CTH expression from the GTex portal. (g-j) Correlation analysis between the CTH and TF transcripts in healthy human liver tissues using the data collected from GEO Dataset. The statistical analysis for the correlation plots was carried out using Spearman correlation. P<0.05 indicates statistically significant correlations

### Role of Sp1, CREB, and NFY in the transcriptional regulation of human CTH gene

To ascertain the role of these *in silico*-predicted TFs as transcriptional regulators, we altered their expression and measured the CTH promoter activity and endogenous CTH mRNA levels in Huh7 cells. Upregulation of the expression of CREB/Sp1/NFY-A augmented the CTH promoter activity of CTH in a dose-dependent manner (one-way ANOVA, F = 156.5, P<0.0001; one-way ANOVA, F = 42.59, P<0.0001; one-way ANOVA, F = 51.11, P<0.0001) (Fig. 4 a-c). The endogenous CTH mRNA levels were also augmented with increasing levels of -CREB/Sp1/NFY-A (one-way ANOVA, F = 556.9, P<0.0001; one-way ANOVA, F = 26.41, P = 0.0002; one-way ANOVA, F = 23.66, P<0.0001) Fig. 4 a-c). Conversely, co-transfection of CREB and NFY dominant-negative expression plasmids (viz. KCREB and NFYDN), or co-transfection of Sp1 shRNA expression plasmid, resulted in regressed promoter activity (one-way ANOVA, F = 186.3, P<0.0001; one-way ANOVA, F = 146.3, P<0.0001; one-way ANOVA, F = 118.7, P<0.0001; respectively) (Fig. 4 d-f). Consistent with these exogenously transfected CTH promoter-reporter activities, the corresponding endogenous CTH mRNA levels in KCREB, Sp1 shRNA, and NFYDN transfected cells were also diminished (one-way ANOVA, F = 4.154, P = 0.0476; one-way ANOVA, F = 7.192, P = 0.0117; one-way ANOVA, F = 5.478, P = 0.0243; respectively) (Fig. 4 d-f). Thus, these results indicate pivotal role for CREB, Sp1 and NFY in activating the human CTH expression under basal conditions.

**Fig. 4.**
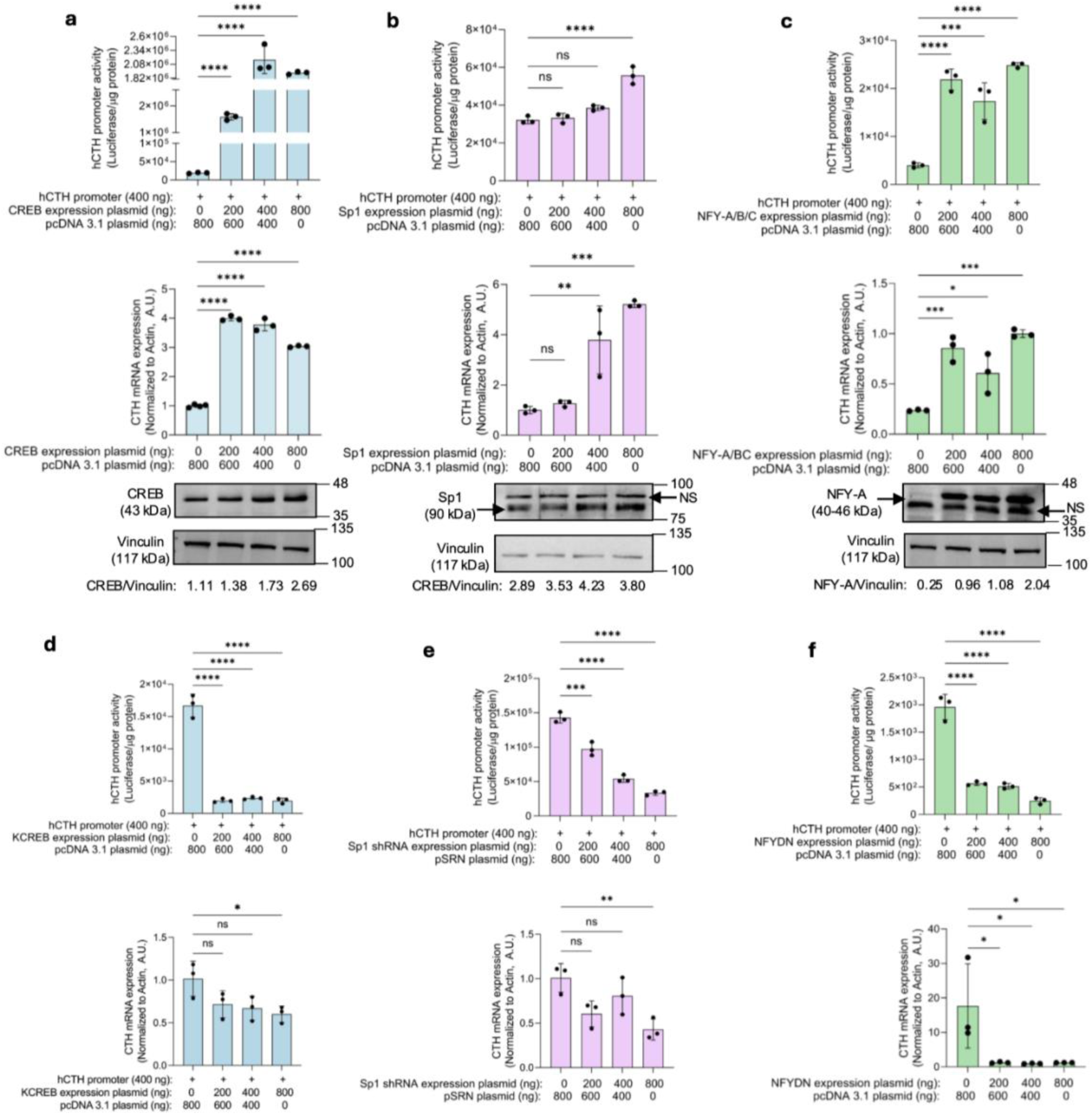
Role of CREB, Sp1 and NFY on the promoter activity and mRNA levels of human CTH gene. CTH pro-275 promoter-luciferase reporter construct was co-transfected with increasing concentrations of CREB, Sp1, and NFY-A/B/C expression plasmids in Huh7 cells. (a-c) CTH promoter activity (top panels) and endogenous CTH mRNA levels (middle panels) increased with increasing CREB, Sp1, and NFY-A protein levels (bottom panels). CTH promoter-reporter construct was co-transfected with increasing concentrations of KCREB, Sp1 shRNA, and NFY-DN expression plasmids in Huh7 cells. (d-f) CTH promoter activity (top panels) and endogenous CTH mRNA decreased (bottom panels) with increasing concentrations of KCREB, Sp1 shRNA or NFYDN expression plasmids. The values in the graphs are mean ± S.D of n = 3 technical replicates. The transfection experiments were repeated at least three times and representative data are shown. The statistical significance was determined by one-way ANOVA with Tukey’s multiple correction as a post-hoc test. ****P<0.0001, ***P<0.001, **P<0.01, *P<0.05, ns: not significant. NS: non-specific protein band

### Effect of mutations in the binding sites of CREB, Sp1, and NFY on human CTH promoter activity

To validate the roles of CREB, Sp1, and NFY in the human CTH gene regulation, we mutated the predicted binding sites by replacing two or more nucleotides such that the binding of these TFs (as per *in silico* predictions) with the CTH promoter was abrogated. Transfection of Huh7 cells with the mutant and wild-type promoter constructs showed significant reductions in the promoter activity in case of the mutants (viz. CREB: ∼0.60-fold, P = 0.0003, Sp1: ∼0.68-fold, P<0.0001, NFYA: ∼0.60-fold, P = 0.0038) (Fig. 5 a-c). Further, because the binding sites of these CREB, Sp1 and NFY-A were adjacent to each other, we hypothesized that these TFs could activate the promoter in a synergistic manner. Hence, CTH-WT promoter (hCTH-pro-275) was co-transfected with a combination of two TFs at a time (CREB and Sp1, CREB and NFY, Sp1 and NFY). Highest promoter activities were observed when either Sp1 or NFY-A were co-expressed with CREB (viz. Sp1 and CREB = 48.8-fold higher as compared to Sp1 alone, P = 0.0024 and (NFY-A and CREB = 44.4-fold higher than compared to NFY alone, P = 0.003) (Fig. 5 d). However, upon co-expression of Sp1 and NFY, the promoter activity only increased by ∼1.34-fold and ∼1.12-fold as compared to Sp1 and NFY, respectively (Fig. 5 d). These results demonstrate that the predicted CREB, Sp1, and NFY-A binding sites in the human CTH promoter are functionally active and constitute transcription activation regulatory elements.

**Fig. 5.**
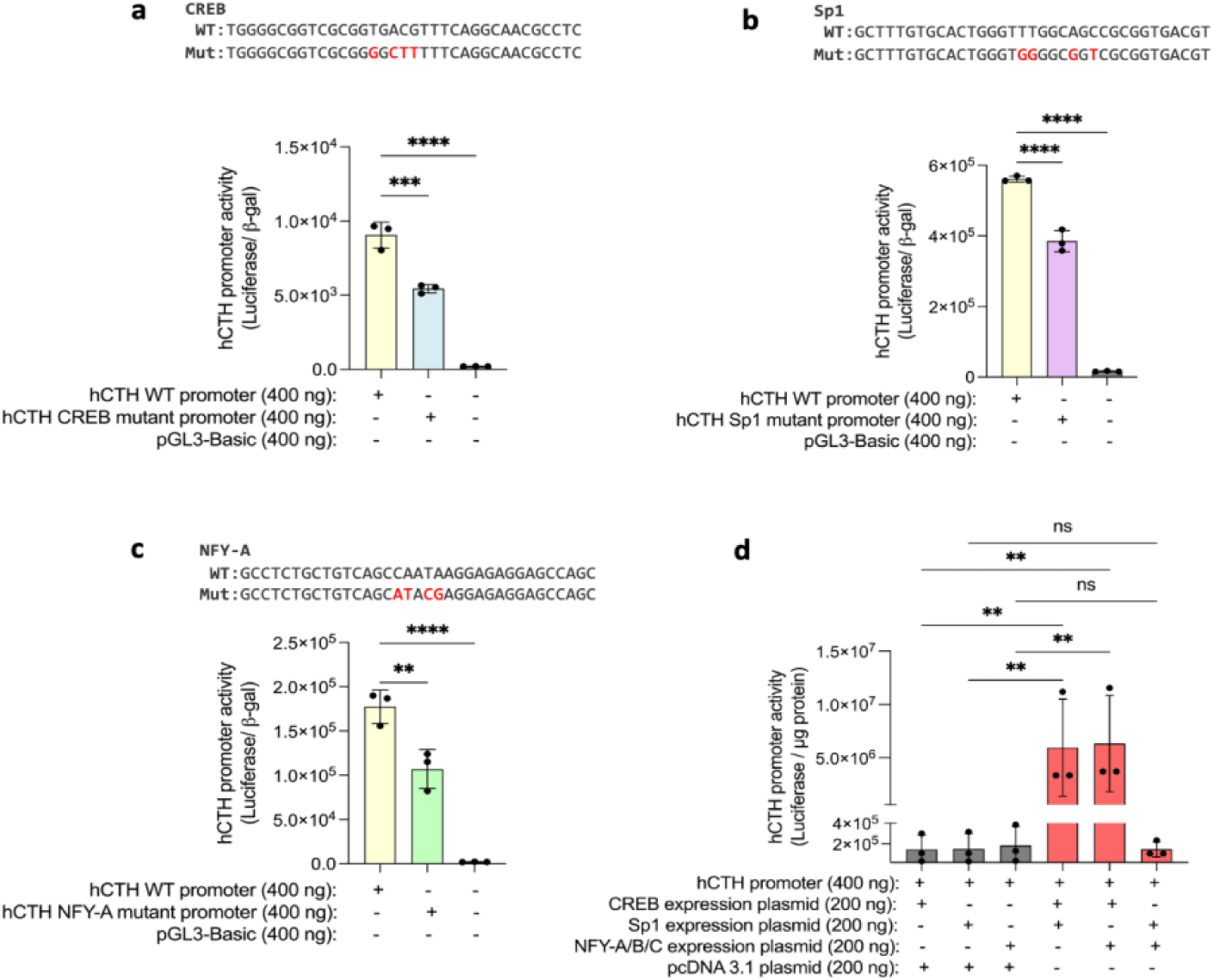
Functional validation of CREB/Sp1/NFY-A binding sites in the human CTH promoter. Huh 7 cells were transfected with core CTH promoter promoter-reporter constructs (WT and mutant) along with β-galactosidase expression plasmid. (a-c) CTH promoter activity upon mutation of the binding sites of CREB, Sp1, and NFY-A in the core promoter region. The mutated bases in CTH promoter are highlighted in red above the plots. Synergistic activity of these TFs was determined by co-transfection of CTH-WT promoter-reporter construct with a combination of two TFs expression plasmids at a time in Huh7 cells. (d) CTH core promoter (WT) activity upon co-overexpression of two TFs. The values in the graphs are mean ± S.D of n = 3 technical replicates. The transfection experiments were repeated at least three times and representative data are shown. The statistical significance was determined by one-way ANOVA with Tukey’s multiple correction as a post-hoc test. ****P<0.0001, ***P<0.001, **P<0.01, ns: not significant

### Confirmation of binding of CREB, Sp1, and NFY-A to endogenous human CTH promoter by chromatin immunoprecipitation (ChIP) assays

To investigate whether the TFs CREB, Sp1, and NFY-A indeed interact with core human CTH promoter region in the context of chromatin (Fig. 3 b) ChIP assays were performed. Antibodies specific to CREB, Sp1 and NFY-A were used to pull-down the DNA-TF complex and the enrichment of CTH promoter was quantified by RT-PCR using primer pairs encompassing the predicted binding sites of the TFs (Fig. 6 a). For all the TFs, under basal/normoxic condition, the enrichment of the CTH promoter was significantly higher in case of TF specific IP than the respective IgG pulled chromatin (negative control) (CREB ∼3 fold, P = 0.0005; Sp1∼7 fold, P = 0.0023; and NFY-A∼14 fold, P = 0.0003) (Fig. 6 b-d), suggesting specificity of pulled-down chromatin being bound to CREB/Sp1/NFY-A and proving the interaction of these TFs with CTH promoter in the predicted region (Fig. 6 a). During ischemia, the enrichment of CTH promoter in Sp1-specific IP was significantly lesser than normoxia (∼4-fold, P = 0.003) (Fig. 6 b), whereas no difference in the enrichment was found for CREB and NFY-A specific IP (Fig. 6 c and d). During reperfusion, significantly higher enrichment as compared to normoxia was observed for CREB (∼13-fold, P<0.0001) (Fig. 6 c). In contrast to this, the enrichment of CTH promoter in NFY-A specific IP was significantly diminished (∼10-fold, P<0.0001) than normoxia or ischemia (Fig. 6 d). While the enrichment of CTH promoter for Sp1-specific IP was statistically higher than ischemia (∼3.5-fold, P = 0.01), no significant difference was observed between normoxia and reperfusion conditions (Fig. 6 b). The differential enrichment of the promoter indicates differential binding of the TFs to the promoter. Hence, we further investigated if differential binding of the TFs to CTH promoter was a consequence of altered endogenous protein levels of respective TFs. Surprisingly, no difference in the protein levels of CREB and Sp1 was found (Fig. 6 e, g, and h). While the protein level of NFY-A was approximately 3-fold lesser (P = 0.05) in ischemia and reperfusion samples as compared to normoxia (Fig. 6 f and i), it did not correlate with CTH promoter enrichment under normoxia, ischemia or reperfusion. These findings suggest that the dysregulation of CTH during ischemia and reperfusion may not be completely associated with transcriptional activity of the studied TFs.

**Fig. 6.**
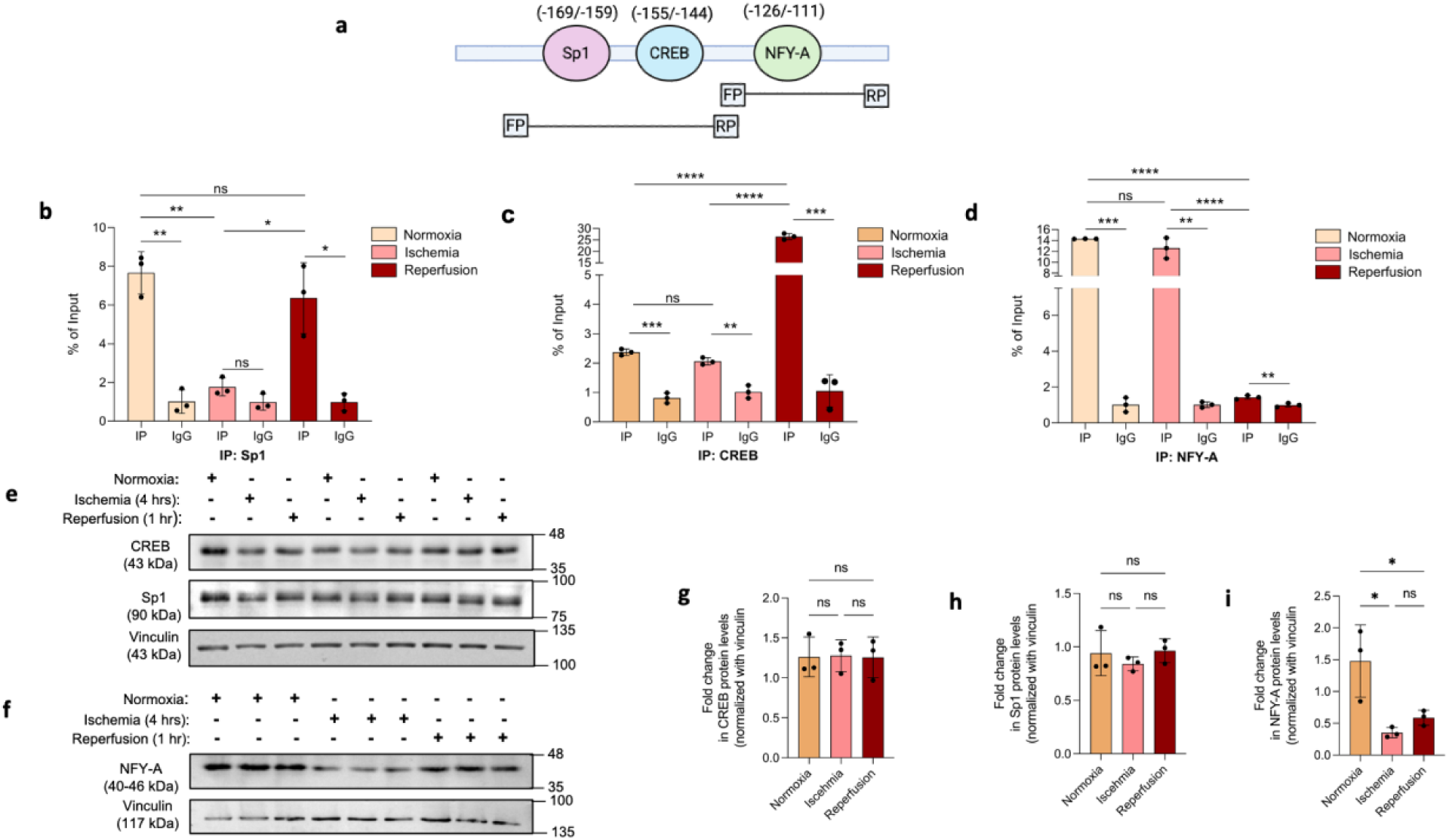
Evidence of interactions of CREB, Sp1, and NFY-A with human CTH promoter in the context of chromatin. ChIP assay followed by RT-PCR was performed to test the interactions of CREB, Sp1, and NFY-A with the human CTH promoter. (a) Enrichment of CTH promoter was measured using primer pairs spanning the predicted binding sites of each TF in the core promoter domain. (b-d) Enrichment of CTH promoter domains after ChIP using antibodies specific to CREB, Sp1, and NFY-A under normoxia, ischemia and reperfusion conditions. (e-i) Endogenous protein levels of TFs under normoxia, ischemia and reperfusion conditions in Huh7 cells, and respective densitometry plots. The values in the graphs of panel b-d are represented as mean ± SD of n = 3 technical replicates. The values in the graphs of panel g-i are represented as mean ± SD of n = 3 biological replicates. Unpaired *t*-test or one way ANOVA with Brown-Forsythe multiple correction as a post-hoc test was used for statistical analysis. ****P<0.0001, ***P<0.001, **P<0.01, *P<0.05, ns: not significant

### Post-transcriptional regulation of CTH gene by miRNAs under basal condition

MicroRNAs are crucial post-transcriptional regulators and play a significant role in regulating the protein abundance by either degrading the target mRNA or by inhibiting their translation. In order to identify novel CTH-targeting miRNAs, we performed rigorous *in silico* analysis using miRNA prediction tools and databases in a hierarchical approach resulting in the selection of hsa-miR-150-5p and miR-510-5p as potential miRNAs that could target human CTH (Fig. 7 a, Fig. S3 a and b). The binding sites for these miRNAs were also conserved in the 3′-UTR of mouse and rat CTH (Fig. S3 c). To test their regulatory role, CTH 3′-UTR reporter construct was co-transfected with increasing doses of miR-150-5p or miR-510-5p overexpression plasmids in Huh7 cells. A dose-dependent decrease in the 3′-UTR -reporter activity (miR-150-5p: one way ANOVA, F = 102.5, P<0.0001; miR-510-5p: one way ANOVA, F = 10.69, P = 0.0036) as well as endogenous CTH protein was observed (Fig. 7 b and c). The overexpression of both miRNAs was confirmed by RT-PCR (miR-150-5p: one way ANOVA, F = 38.51, P<0.0001; miR-510-5p: one way ANOVA, F=1030, P<0.0001) (Fig. S3 d and e). Conversely, specific miRNA inhibitors against miR-150-5p and miR-510-5p were used to knockdown their expression in Huh7 cells. These miRNA inhibitors increased the endogenous CTH mRNA (Fig. S3 f and g) as well as protein levels (Fig. 7 d and e), validating CTH as a target of miR-150-5p and miR-510-5p. The decrease in expression of miR-150-5p and miR-510-5p in respective samples was confirmed by RT-PCR (Fig. S3, h and i).

**Fig. 7.**
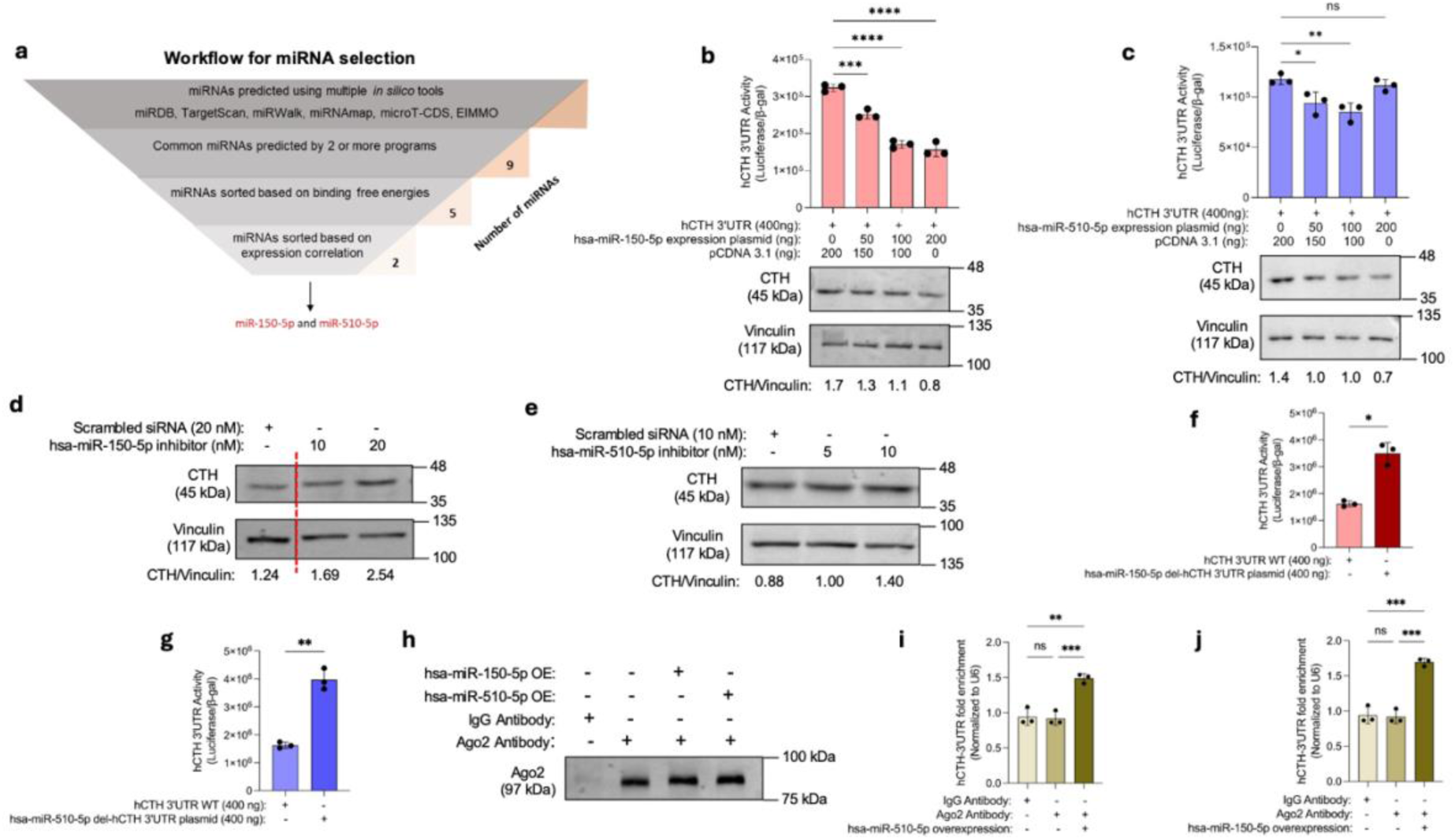
Direct interactions of the microRNAs hsa-miR-150-5p and hsa-miR-510-5p with human CTH 3′-UTR. (a) Schematic showing multiple factors that were considered to shortlist hsa-miR-150-5p and hsa-miR-510-5p as candidate miRNAs that could target human CTH gene. (b and c) Co-transfection of human CTH 3′-UTR constructs with increasing concentrations of either miR-150-5p or miR-510-5p expression plasmids in Huh7 cells resulted in decreased reporter activity and endogenous CTH protein in a concentration-dependent manner. (d and e) Inhibition of endogenous miR-150-5p and miR-510-5p using respective miRNA inhibitors decreased endogenous miRNA levels and increased endogenous CTH protein levels. (f and g) Deletion of miRNA binding sites on human CTH 3′-UTR enhanced its activity. (i and j) RT-PCR followed by Ago2 IP reveals CTH mRNA was highly enriched in miR-150-5p and miR-510-5p overexpressed samples. The luciferase reporter activity was normalized with β-galactosidase activity and the values obtained for each construct were represented as hCTH 3′-UTR activity. All the values in the graphs are represented as mean ± S.D of n = 3 technical replicates. The transfection experiments were repeated at least three times and representative data are shown. Unpaired *t*-test or one-way ANOVA with Tukey’s multiple correction as a post-hoc test was used for statistical analysis. ****P<0.0001, ***P<0.001, **P<0.01, *P<0.05, ns: not significant

### Confirmation of the interactions of miR-150-5p and miR-510-5p with CTH 3′-UTR

To probe whether miR-150-5p and miR-510-5p indeed interact with the human CTH 3′-UTR in the predicted region, we abrogated the binding sites of these miRNAs in the CTH 3′-UTR -luciferase reporter construct. Deletion of the binding sites of either of the miRNAs increased the 3′-UTR reporter activity by at least 2-fold (miR-150-5p del, P = 0.0109; miR-510-5p del, P = 0.0054) (Fig. 7, f and g). We also verified the interaction of these miRNAs with CTH 3′-UTR by Ago2 immunoprecipitation (Ago2 IP) followed by assessment of enriched CTH mRNA by RT-PCR because Ago2 protein is a component of RNA-induced silence complex assembly. The specificity of Ago2 IP was confirmed using western blotting (Fig. 7 h). The enrichment of CTH mRNA was significantly higher in samples overexpressing either miR-150-5p (∼1.5-fold, P = 0.0001) or miR-510-5p (∼1.5-fold, P=0.001) compared untreated control (Fig. 7 i and j), providing evidence for interactions of both these miRNAs with CTH 3′-UTR. Overall, these results confirm that miR-150-5p and miR-510-5p down-regulate CTH expression by interacting with the 3′-UTR of CTH mRNA at the designated binding sites (Fig. S3 b).

### Role of miR-150-5p and miR-510-5p in regulating CTH expression in hepatocytes during ischemia-reperfusion

Having established CTH as a regulatory target of miR-150-5p and miR-510-5p under basal conditions, we aimed to explore their roles in regulating CTH expression during ischemia and reperfusion. Accordingly, we quantified the miR-150-5p and miR-510-5p expression levels in normoxia, ischemia and reperfusion conditions. The expression of both miRNAs peaked after 4 hours of ischemia (miR-150-5p, ∼7 fold as compared to normoxia, P = 0.0004; miR-510-5p, ∼2 fold as compared to normoxia, P<0.0001), followed by a sharp decrease after 1 hr of reperfusion (miR-150-5p, ∼7 fold with respect to ischemia, P = 0.0005; miR-510-5p, ∼4-fold with respect to ischemia, P<0.0001). The expression levels of both miRNAs were negatively correlated with CTH expression in respective conditions (Fig. 8 a-f).

**Fig. 8.**
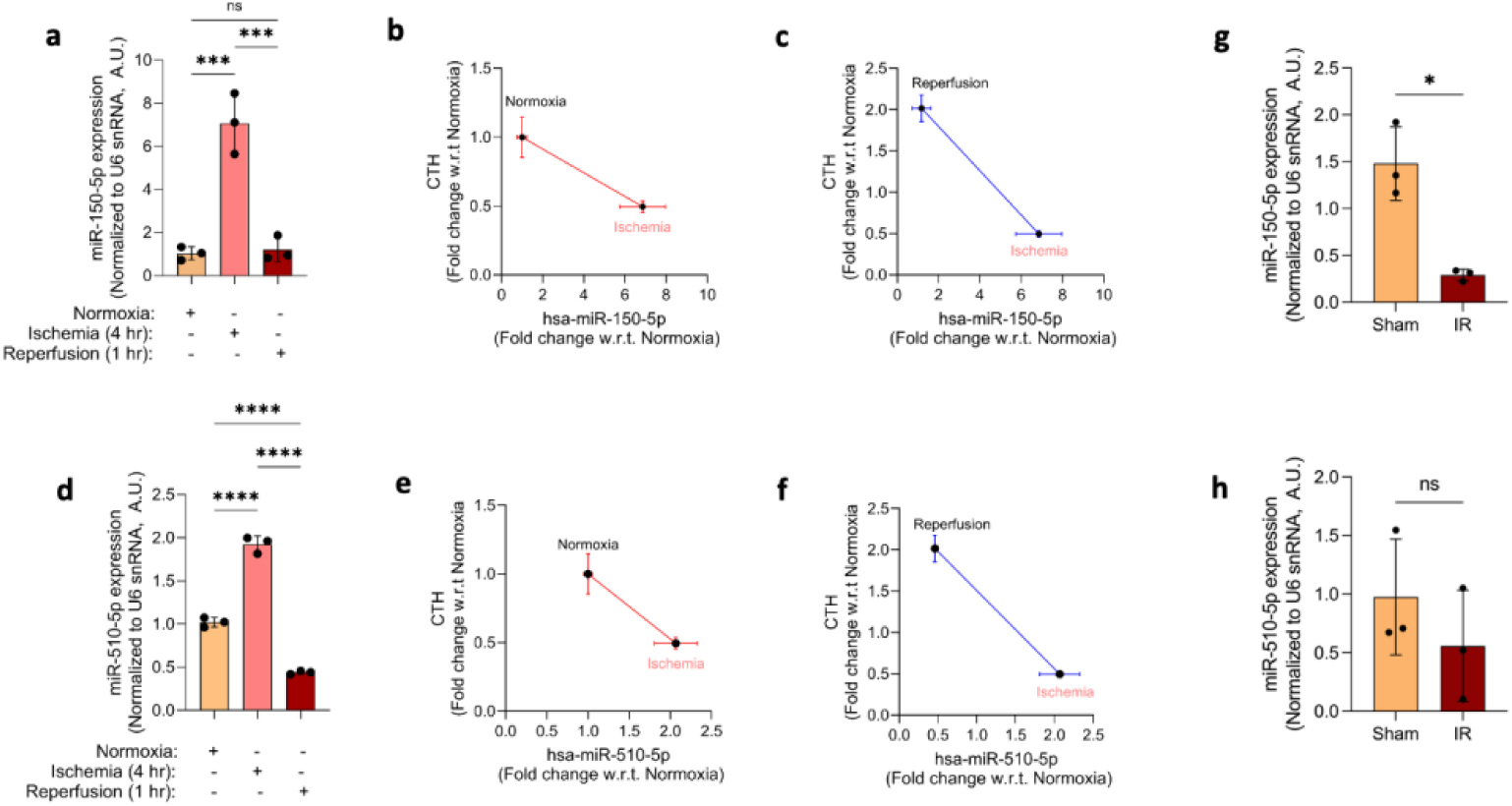
miR-150-5p and miR-510-5p post-transcriptionally regulate CTH expression under ischemia and reperfusion conditions. Huh7 cells were subjected to normoxia, ischemia (4 hr), and reperfusion (1 hr) and microRNA expression levels were measured using RT-PCR. (a) Expression levels of miR-150-5p in Huh7 cells in normoxia, ischemia and reperfusion conditions. (b and c) Correlation between miR-150-5p and CTH expression during normoxia/ischemia conditions and ischemia/reperfusion in Huh7 cells. (d) Expression levels of miR-150-5p in Huh7 cells in normoxia, ischemia and reperfusion conditions. (e and f) Correlation between miR-510-5p and CTH expression during normoxia/ischemia conditions and ischemia/reperfusion in Huh7 cells. (g and h) Expression levels of miR-150-5p and miR-510-5p in mice IR liver (n = 3). All the values in the graphs (except panel g and h) are represented as mean ± S.D of n = 3 technical replicates. The *in vitro* IR experiments were repeated at least three times and representative data are shown. Unpaired t-test or one-way ANOVA with Tukey’s multiple correction as a post-hoc test was used for statistical analysis. ****P<0.0001, ***P<0.001, *P<0.05, ns: not significant

Since the binding sites for miR-150-5p and miR-510-5p are conserved between human and mouse CTH 3′-UTR sequences, we quantified the expression of these miRNAs in control and IR mouse liver tissues. The expression of miR-150-5p (but not that of miR-510-5p) was significantly diminished in IR mouse liver samples (∼7-fold, P = 0.0319) with respect to control) (Fig. 8 g and h) while the CTH protein levels were augmented (∼1.8-fold, P= 0.0096 with respect to control) (Fig. 1 e). Altogether, these results highlight the crucial roles of miR-150-5p and miR-510-5p in the post-transcriptional regulation of CTH expression under basal/normoxia, ischemia and reperfusion conditions in hepatic cells across species.

### Role of histone modifications in CTH regulation under normoxia, ischemia and reperfusion conditions

Ischemia and oxidative stress are known to alter epigenetic signatures in cells by altering DNA methylation or triggering histone modifications [45]. In particular, HDAC6 deficiency in mice has been associated with reduced hepatic injury post-reperfusion [59–61]. Promoter accessibility is a necessary pre-requisite for transcriptional machinery to work, and epigenetic modifications are crucial regulators of DNA accessibility. In view of these findings, we set out to explore whether histone modifications also contribute to CTH dysregulation during IR. Accordingly, we tested the possible roles of commonly studied gene-activating (H3K4 trimethylation aka H3K4me3 and H3K9 acetylation aka H3K9ac) and gene suppressing modifications (H3K9me3 and H3K27me3) and performed ChIP assays. The enrichment of CTH promoter domains under each condition was quantified by qPCR using four primers sets (viz., primer set 1: -146 bp /+1 bp; primer set 2: -197 bp/-77 bp; primer set 3: -275 bp/-178 bp; primer set 4: -472 bp/-257bp; Table S8) designed to amplify a total length of 472 bp human CTH promoter (Fig. S4 a). The percentage enrichment of each promoter h region was measured (Fig. 9 a-c, Fig. S4 b) and the cumulative area under the curve was calculated and plotted as line plot (Fig. 9 d-f, Fig. S4 c) and bar plot (Fig. 9 g-i, Fig S4 d). For H3K4me3, the highest enrichment of the CTH promoter was observed during ischemia, followed by reperfusion and normoxia (Fig. 9 g). In contrast to this, for H3K9ac, the highest enrichment of the promoter was found in normoxia followed by ischemia and reperfusion; however, no significant difference in the enrichment between ischemia and reperfusion was found (Fig. 9 h). For H3K27me3 modification, the enrichment of the promoter was highest in ischemia, whereas for normoxia and reperfusion samples, the enrichment of the promoter was not statistically different from respective negative control (IgG) (Fig. 9 i). Similarly, for H3K9me3, the enrichment of promoters in IP samples of none of the groups was significantly different from their respective negative controls (IgG) (Fig. S4 d). Thus, different histone modifications seem to be involved in the gene regulation of CTH at the epigenetic level.

**Fig. 9.**
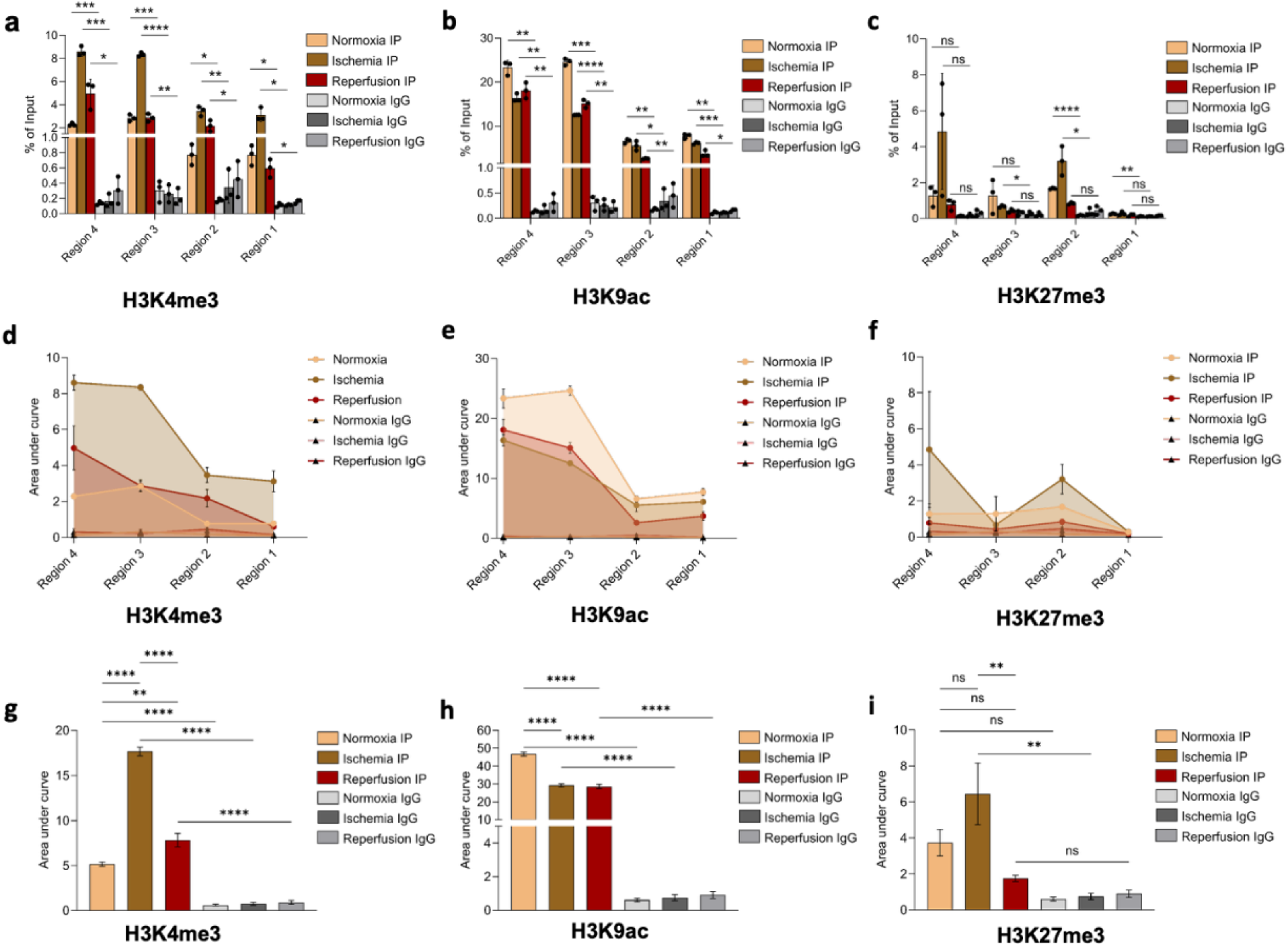
Epigenetic regulation of CTH expression by H3K4me3, H3K9ac, and H3K27me3 modifications during normoxia, ischemia and reperfusion conditions. The CTH promoter DNA was enriched by ChIP for different conditions and was quantified using RT-PCR using four sets of primers that amplified a total of ∼470 bp promoter region. (a-c) The percentage enrichment of all the four regions of CTH promoter for each histone modification and for every treatment condition (a-c). (d-f) Line plots depicting the calculated cumulative area under curve (AUC) for the entire enriched promoter. (g-i) Bar plot depicting cumulative AUC for each histone modification under different treatment conditions. All the values in the graphs are represented as mean ± S.D of n = 3 technical replicates. Unpaired *t*-test or one-way ANOVA with Šidák multiple correction as a post-hoc test was used for statistical analysis. ****P<0.0001, ***P<0.001, **P<0.01, *P<0.05, ns: not significant

## DISCUSSION

### Overview

H_2_S treatment has been widely used in preclinical studies to protect cardiac, renal, hepatic or brain tissues in animal models of hypoxic, ischemic or reperfusion injuries. Moreover, manipulating H_2_S bioavailability by CTH gene knockdown/knockout, or by perturbing gut microbiome, tissue damage caused due to oxidative stress can be prevented. Hence, modulating H_2_S bioavailability by controlling CTH expression in an oxidative milieu could potentially serve as an effective therapeutic approach. The expression of a given gene is not a standalone event but involves a complex interplay between various molecular factors including TFs, miRNAs and epigenetics. In this study, we report the role of novel TFs, miRNAs, and critical epigenetic factors in modulating human CTH expression under basal as well as during ischemia and reperfusion pathophysiology.

### Molecular mechanisms of transcriptional regulation of CTH gene

Previously Yin et. al, reported the core promoter of CTH lies within the -592/+139 region and that harboured four Sp1 binding sites [62]. In this study, using stringent computational analysis and promoter deletion constructs, we further narrowed down the core promoter of human CTH within -275 bp form TSS, which, besides Sp1, also harbors binding sites for CREB, and NFY-A (Fig. 3). It is well-established that Sp1 binds to the GC/GT box, CREB binds to the CRE and NFY binds to the CAAT box in promoter of various genes (reference). These elements are important eukaryotic promoter domains, and these cis-regulatory motifs are also conserved in the mouse CTH promoter, indicating their key roles in CTH expression. Indeed, over-expression of Sp1, CREB, and NFY-A could significantly upregulate CTH promoter activity under basal/unstimulated conditions (Fig. 4). In corroboration, mutation of these TF binding sites in the CTH promoter and ChIP assays confirmed their role as transcriptional activators under physiological conditions in hepatic cells. Not only do these TFs independently augment CTH promoter activity, but they also act in a synergistic manner (Fig. 5). The synergistic activity can be explained by the close proximity of their binding sites CTH promoter. This is consistent with previous reports documenting close proximity of Sp1, CREB and NFY binding sites in the promoters of several eukaryotic genes [46, 63, 64]. Besides, the proximity of the binding sites, physical interactions between CREB and Sp1, and CREB and NFY-A have also been previously reported [65, 66]. Nonetheless, this is the first study that reports CREB and NFY as novel transcriptional activators for human CTH and that these TFs may act in a coordinated manner to activate CTH gene expression.

Moreover, the transcription factors Sp1, CREB and NFY-A are involved in the regulation of stress-responsive genes including antioxidant genes [67–73]. Several studies have reported CTH dysregulation in ischemia-reperfused tissues, oxidative stress and calorie or amino acid restriction models. IR is a condition with immense stress driven by increased reactive oxygen species (ROS) and transient absence of nutrients. Hence, we asked whether these TFs could explain the CTH dysregulation during I/R? Of note, Sp1 or CREB protein abundance did not change significantly during I/R. While the NFY-A protein levels were diminished during ischemia and reperfusion, it did not mirror CTH expression during these conditions (Fig. 6). Furthermore, the binding of these TFs to the CTH promoter during these conditions did not mirror their protein abundance (Fig. 6).

A possible explanation for these observations could be that post-translational modifications of these TFs under excess ROS (due to IR stress) might result in their altered DNA binding activity. This speculation is supported by a previous report, wherein reduced binding of Sp1 to CBS promoter after 2 hr of ischemia and 6 hr of reperfusion was attributed to its phosphorylation [74]. Similarly, ROS induced phosphorylation of CREB at Ser133, resulted in its activation, nuclear translocation and enhanced DNA binding activity [75]. Furthermore, CREB activation has been previously used to ameliorate the liver damage caused by IR in mice [76]. On the contrary, the genome-wide occupancy of NFY reduced under oxidative stress conditions, suggesting reduced DNA binding activity [67]. The reduced DNA binding activity under elevated oxidative stress was attributed to oxidation of cysteine residues of NFY-B subunit, which prevents its interaction with NFY-C and thereby abrogating trimerization of NFY-A, NFY-B and NFY-C subunits-an event crucial for the activity of NFY [77, 78]. Although we have not carried out experiments in this study to test these post-translational modifications, our ChIP assay findings (Fig. 6) are in agreement with these explanations.

### Molecular mechanisms of regulation of CTH at post-transcriptional level: roles of microRNAs

Several miRNAs have also been reported to regulate CTH expression under diverse pathological conditions, including members of the miR-30 family in cardiac ischemic injury [27] miR-21 and miR-143 during smooth muscle cell differentiation [28, 29], miR-27b in atherosclerosis [30], miR-133a in cardiac hypertrophy [31], and miR-186 in proinflammatory THP-1 macrophages [33]. Epigenetic mark such as DNA methylation has also been reported to regulate CTH in expression in FOXC1 dependent manner [32]. Nevertheless, the mechanism of regulation of CTH gene at the post-transcriptional level during IR is poorly understood. This study aimed to identify miRNAs that directly target CTH gene under basal and IR conditions. Our rigorous computational analysis followed by experimental validation identified miR-150-5p and miR-510-5p as novel post-transcriptional regulators of human CTH under basal and IR conditions (Fig. 7 and 8). As the binding site of these miRNAs in the 3′-UTR of CTH is conserved across humans and mice, we observed an inverse correlation between miR-150-5p levels and CTH expression in reperfused mice liver (Fig. 1 and 8). This is consistent with the previous report demonstrating significantly reduced miR-150-5p levels in transplanted human liver tissue [79]. This concordance between our findings and previous clinical observation suggests a clinical relevance of miR-150-5p in the pathophysiology associated with liver IR.

### Role of histone modifications in CTH gene regulation

Epigenetic regulation of CTH during IR appears to be highly coordinated, rather than being governed by a simple activation or inhibition mechanism. In our study, both H3K4me3 and H3K27me3 and, a canonical activation and repressive mark, respectively, were enriched during ischemia (Fig. 9). Contrary to this, H3K9ac enrichment markedly reduced during ischemia (Fig. 9 h). With reperfusion, enrichment of H3K4me3 and H3K27me3 diminished but H3K9ac remained unchanged with simultaneous recovery of CTH expression (Fig. 9). The increase in H3K27me3 during ischemia is consistent with previous observations in rat hippocampal tissue, where ischemia induced H3K27me3 accumulation followed by a time-dependent decline during reperfusion [80]. Similarly, safeguarding acetylation footprints by inhibiting their erasers-HDACs, for instance HDAC3, was sufficient to upregulate CTH expression and H_2_S production in vascular smooth muscle cells [81]. In fact, inhibition of HDACs is a proven strategy to confer protection against ischemic insult [82–86]. The abundance of H3K4me3 modification at the CTH promoter, however, indicates another layer of complexity due to its widely accepted role as an activator but inverse association with CTH expression observed in this study. Due to the coexistence of H3K4me3 with the repressive H3K27me3 mark, a mutually exclusive event, it may be assumed that CTH promoters exhibit bivalent behavior.

Bivalent chromatin states have been reported previously for developmental genes [87–90]. A recent characteristic feature of bivalent chromatin is that these regions of DNA are CpG rich and bivalency protects it from hypermethylation, thereby maintaining the genes in a poised state (repressed transiently but can be rapidly activated) [91]. The presence of multiple CpG sites in human CTH promoter (Fig. S4 e) make the simultaneous presence of H3K4me3 and H3K27me3 relevant, when the dynamic of CTH expression during basal, ischemia, and reperfusion is considered. Methylation dependent repression of CTH has been previously reported in mouse macrophages [92]. Similarly, methylation of CTH promoter in kidney cell line where IR was associated with increased methylation of the CTH promoter and reduced CTH expression, whereas inhibition of DNA methylation during ischemia restored CTH expression [93]. Loss of promoter bivalency was associated with increased DNA methylation and sustained transcriptional repression in cancer cells [94]. Besides, we also observed that multiple histone methylation marks are elevated during ischemia despite their canonical chromatin accessibility roles-reflecting a broader consequence of ischemic cellular milieu. Ischemia is accompanied with heightened oxidative stress and metabolic perturbations which may influence the activity of histone demethylases [95–98]. Moreover, oxidative stress, and exposure to heavy metals have been reported to alter demethylase activity and increase global levels of H3K9me2, H3K4me3, and H3K27me3 [99–101]. Taken together, it appears that ischemia induces a globally methylated and transcriptionally constrained CTH promoter, while it retains a bivalent state. Reperfusion reverses the inactive epigenetic state by reducing the inactivation mark (H3K27me3) without altering the activation epigenetic marker (H3K9ac), which is accompanied by increased CTH expression. Because of the key role of CTH in maintaining metabolic and redox homeostasis, such dynamic interplay between histone methylation, promoter bivalency, could be particularly important to prevent CTH from undergoing long-term repression during ischemia and permitting rapid recovery during reperfusion.

## Conclusions

Our investigations identified three TFs (Sp1, CREB and NFY-A), two miRNAs (miR-150-5p and miR-510-5p), and three histone modifications (H3K4me3, H3K27me3, and H3K9ac) that regulate human CTH gene expression at the transcriptional, post-transcriptional and epigenetic levels, respectively. To our knowledge, this is the first report that unravels molecular mechanism of CTH regulation during ischemia and reperfusion as independent conditions. In summary, ischemia causes reduced binding of Sp1 on CTH promoter, increased levels of miR-150-5p and miR-510-5p, and reduced H3K9ac, together resulting in reduced CTH expression. On the contrary, reperfusion causes increased Sp1 and CREB binding on CTH promoter, reduced miR-150-5p and miR-510-5p expression, reduced H3K27me3, thereby restoring the CTH expression. The plausible molecular mechanisms of human CTH gene regulation and molecular factors molecular factors involved are illustrated in Fig. 10.

**Fig. 10.**
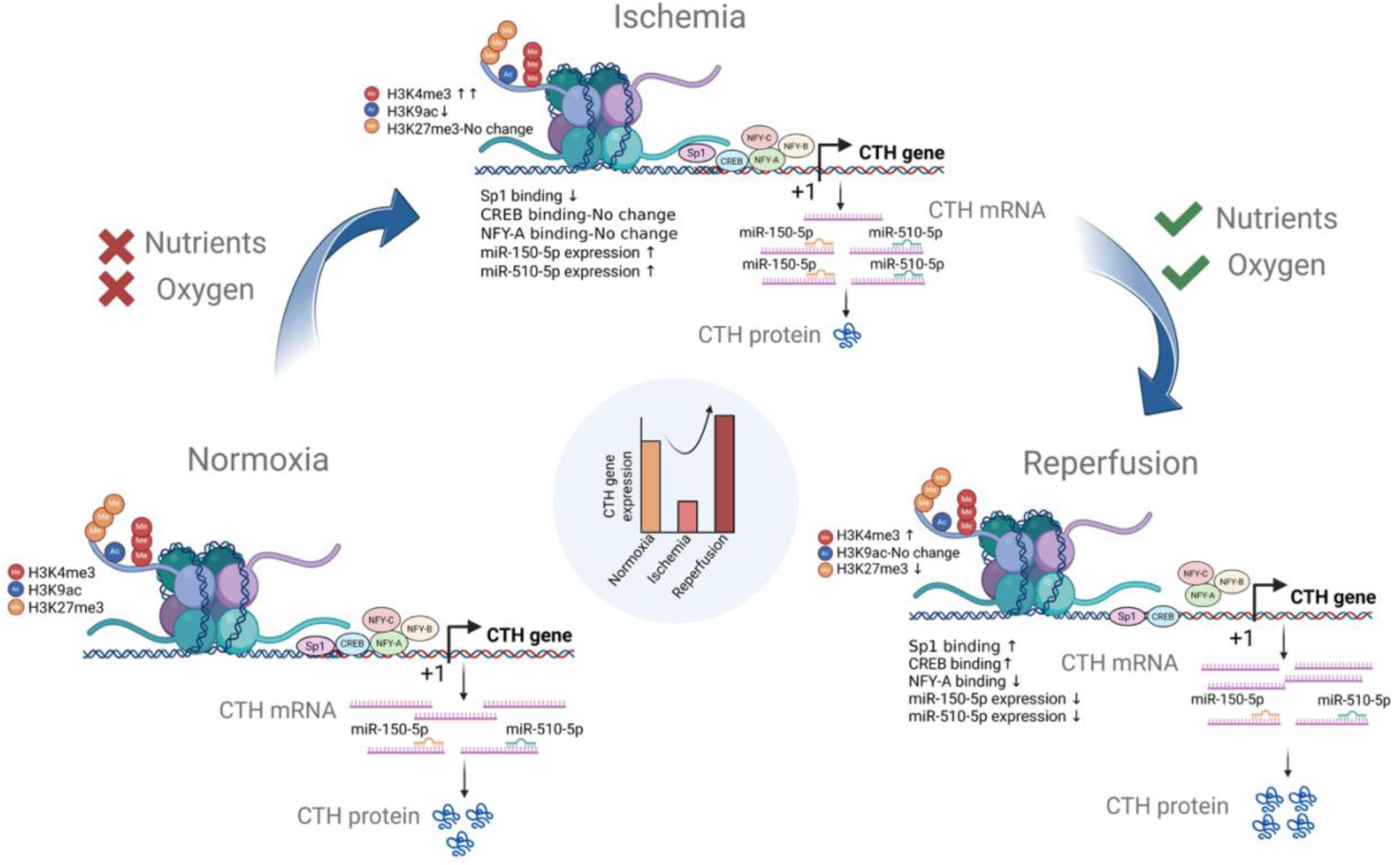
Plausible mechanism of CTH regulation under normoxia, ischemia, and reperfusion conditions. Under normoxia condition, the acetylation of H3K9 and methylation of H3K4 renders the DNA encompassing CTH promoter accessible to the transcription factors Sp1, NFY, CREB, resulting in the transcriptional activation of CTH expression. Post-transcriptionally, miR-150-5p and miR-510-5p binding to CTH mRNA in the 3′-UTR regions, causes mRNA degradation. Equilibrium between the transcriptional and post-transcriptional regulation of CTH results in the baseline expression of CTH. However, during ischemia, H3K9 is deacetylated, methylation of H3K4 dramatically increases. Consequently, CREB and NFY binding on CTH promoter remains unaltered while Sp1 recruitment is drastically reduced. The levels of miR-150-5p and miR-510-5p are augmented. Collectively, these events suppress CTH mRNA/protein expression. On the contrary, during the reperfusion phase, methylation of H3K27 and H3K4 are diminished, acetylation of H3K9 remains unaltered. Binding of Sp1 to the CTH promoter (which is reduced during ischemia) is augmented. Likewise, CREB binds with a higher affinity to the CTH promoter (when compared to normoxia or ischemia). NFY recruitment is significantly diminished. Both miR-150-5p and miR-510-5p levels return to baseline levels. Overall, these events augment CTH expression during reperfusion

## Competing Interests

The authors declare that they have no competing interests.

## Author Contributions

Anand Kumar Patel designed the study, performed experiments, and wrote the manuscript. Navasuja Ganesh, Vatsal Arya, and Gopikrishna Rathinavel performed experiments. Piyushkumar R. Kapopara performed experiments and contributed to drafting the manuscript. Vikas Arige assisted with experiments and contributed to drafting the manuscript. Nitish R. Mahapatra contributed to study design, supervised the research, acquired funding, and reviewed and edited the manuscript.

## Data availability

All data needed to evaluate the conclusions in this article are available in the methods and/or the supplementary material of this article. Any additional information on the findings of this study is available on request from the corresponding author.

## Ethics Approval

This study was performed in line with the principles of the Declaration of Helsinki. Approval was granted by the Institute Animal Ethics Committee of Indian Institute of Technology Madras, Chennai, India (Approval number: IITM/IAEC/NRM-2/05/2023 dated May 27, 2023).

## Acknowledgements

The authors acknowledge the research fellowships provided by Indian Institute of Technology Madras and the Ministry of Education, Government of India to Anand Kumar Patel, Navasuja Ganesh, Piyushkumar R. Kapopara, Vatsal Arya, Gopikrishna Rathinavel and Vikas Arige. The authors also thank Dr. Anupama Vijaykumar, Cardiovascular Genetics laboratory, IIT Madras for her valuable suggestions for improving the manuscript.

## Funding

The project was funded by Board of Research in Nuclear Sciences, Government of India (grant number: 2011/37B/48/BRNS/2026). This study was also supported by grants from the Department of Biotechnology (BT/PR54616/BMS/85/484/2024), Indian Council of Medical Research (EM DR/SG/13/2023-5904) and Anusandhan National Research Foundation (CRG/2023/005900), Government of India.

## Abbreviations

CTH: Cystathionine gamma-lyase
CBS: Cystathionine beta-synthase
H_2_S: Hydrogen sulfide
DEN: Diethylnitrosamine
CCl_4_: Carbon tetrachloride
IRI: Ischemia Reperfusion Injury
3’-UTR: 3’-Untranslated Region
NCBI: National Center for Biotechnology Information
GTEx: Genotype Tissue Expression
CREB: c-Amp Response Element Binding protein
NFY: Nuclear Factor Y
NFY-A: Nuclear Factor Y Subunit Alfa
NFY-B: Nuclear Factor Y Subunit Beta
NFY-C: Nuclear Factor Y Subunit Gamma
Sp1: Specificity protein 1
AP-2α: Transcription Factor Activator Protein 2 alpha
ChIP: Chromatin Immunoprecipitation

## Notes

### Competing Interest Statement

The authors have declared no competing interest.

### Summary of Updates

Results section-CTH expression is dysregulated during ischemia and reperfusion updated to cite Supplementary Table S11; Results section-Putative TF binding sites in the human CTH promoter and correlation analysis between the transcript levels of the predicted TFs and CTH across human tissues updated to reflect the renumbering of Supplementary Tables S11 and S12 as S12 and S13, respectively; new Supplementary Table S11 containing CTH mRNA expression data from human liver tissues (GSE151648) added; miRNA prediction tool citations moved from the Supplementary Information to the Materials and Methods section; Figure 7E labeling updated; Figure 10 summary schematic revised; authorship order updated; declarations and statements moved before the References section.

