## Supplementary Information for "Dynamic regulation of cystathionine gamma-lyase expression during normoxia, ischemia, and reperfusion in liver: interplay of transcription factors, microRNAs, and histone modifications"

*by*

Anand Kumar Patel[a], Navasuja Ganesh[a], Vatsal Arya[a], Piyushkumar R. Kapopara[a,c], Gopikrishna Rathinavel[a], Vikas Arige[a,b], Nitish R. Mahapatra[a#]

*from the*

[a]Department of Biotechnology, Bhupat and Jyoti Mehta School of Biosciences, Indian Institute of Technology Madras, Chennai 600036, India

[b] Department of Pharmacology and Physiology, School of Medicine & Dentistry, University of Rochester, New York, USA

[c] TriLink Biotechnologies, part of Maravai Life Sciences, San Diego, California 92122, USA

*Correspondence to:

Dr. Nitish R. Mahapatra, Department of Biotechnology, Bhupat and Jyoti Mehta School of Biosciences, Indian Institute of Technology Madras, Chennai 600036, India.; Tel: 91-44-2257-4128
Orchid ID: 0000-0002-0695-3510

#### Supplementary Tables

**Table S1. List of primers used to prepare human CTH promoter-reporter deletion constructs**

| Primer name | Forward Primer (5′ to 3′) |
|---|---|
| hCTH-pro-816 | CGGGGTACCCGCAGGTGAGTGGTAAACAATCC |
| hCTH-pro-616 | CGGGGTACCGCTGCCAGAGTTGTGCATAGACG |
| hCTH-pro-375 | CGGGGTACCGGCAAAGGATGGTGGTCAGAG |
| hCTH-pro-275 | CGGGGTACCCACGTCTCGGGCGAACC |
| hCTH-pro-123 | CGGGGTACCTTGTGCACTGGGTGGGG |
| hCTH-pro-75 | CGGGGTACCGCTGTCAGCCAATAAGGAGC |
| **Primer name** | **Reverse Primer (5′-3′) (common for all the hCTH-pro forward primers)** |
| hCTH+92 | CCGCTCGAGAAAGCACGCAGGTAGAGCCGAC |

**Table S2. Primers used to clone human CTH 3'UTR**

| Primer name | Forward Primer (5′ to 3′) | Reverse Primer (5′ to 3′) |
|---|---|---|
| hCTH_3'UTR | CCGCTCGAGGCACTGTTATTA TAGCACCCTCCA | CCCAAGCTTAAGCTCCTCCTCCT GGTTTC |

**Table S3. Primers used to clone pre-miR-510**

| Primer name | Forward Primer (5′ to 3′) | Reverse Primer (5′ to 3′) |
|---|---|---|
| hsa-mir-510 | GTCCTGAAAACTGCCATGTGA | ACAGGCAGATAGGAACACCA |

**Table S4. Primers used for side directed mutagenesis of TF binding sites in hCTH-pro-275 plasmid**

| Primer name | Forward Primer (5′ to 3′) | Reverse Primer (5′ to 3′) |
|---|---|---|
| hCTH-CREB-SDM | GGTGGGGCGGTCGCGGGGCTTTTT CAGGCAACGCCTC | GAGGCGTTGCCTGAAAAAGCCCCGCG ACCGCCCCACC |
| hCTH-Sp1-SDM | ACTCCGCTTTGTGCACTGGGTTTG GCAGCCGCGGTGACGT | ACGTCACCGCGGCTGCCAAACCCAGT GCACAAAGCGGAGT |

**Table S5. Primers used to mutate the binding sites of miRNAs in human CTH 3′-UTR**

| Primer name | Forward Primer (5′ to 3′) | Reverse Primer (5′ to 3′) |
|---|---|---|
| CTH_3'UTR_mir-150_ del | CCGCTCGAGGCACTGTTATTATAGC ACCCTCCA | GCCTGTAGTCCCAGCTACTTCG AACCC |
| CTH_3'UTR_mir-510_del | GCTTCCTGTGAAGATCAAATCTTAA TTAAATGGACCAACAATGAGC | GCTCATTGTTGGTCCATTTAATT AAGATTTGATCTTCACAGGAAG C |

**Table S6. List of miRNAs that were predicted to interact with 3′-UTR region of human CTH gene**

| No. | miRNAs | Predicted by number of programs | Binding free energy (kcal/mol) | Correlation with CTH expression |
|---|---|---|---|---|
| 1 | hsa-miR-27b-3p | 6 | -23.6 | Negative |
| 2 | hsa-miR-150-5p | 5 | -22.8 | Negative |
| 3 | hsa-miR-27a-3p | 5 | -20.6 | N/A |
| 4 | hsa-miR-513a-5p | 5 | -27.9 | Positive |
| 5 | hsa-miR-510-5p | 4 | -23.3 | Negative |
| 6 | hsa-miR-450b-5p | 4 | -17.5 | N/A |
| 7 | hsa-miR-936 | 3 | -23.8 | N/A |
| 8 | hsa-miR-1305 | 3 | -18.9 | N/A |
| 9 | hsa-miR-3658 | 3 | -16.8 | N/A |
| 10 | hsa-miR-26b-5p | 2 | -20.8 | N/A |

**Table S7. Primers used for ChIP assays for TFs binding confirmation**

| Primer name | Forward Primer (5′ to 3′) | Reverse Primer (5′ to 3′) |
|---|---|---|
| CTH-275_Sp1_CREB_ChIP | AGGGACTCCAGCTTCACTCC | CCATTGAGCGCAGGCCCGAC |
| CTH-275_NFYA_ ChIP | GACGTTTCAGGCAACGCCTC | AGTGAAGCTGGAGTCCCTGG |

**Table S8: Primers used for ChIP assays for detection of histone modifications**

| Primer name | Forward Primer (5′ to 3′) | Reverse Primer (5′ to 3′) |
|---|---|---|
| CTH-ChIP-Region4 | TTAACCACCCTCCACGTTGTG | CCATGCTGGCTCCTGAGAG |
| CTH-ChIP-Region3 | CCGGCAAAGGATGGTGGTCAGAG | AGTGAAGCTGGAGTCCCTGG |
| CTH-ChIP-Region2 | AGGGACTCCAGCTTCACTCC | CCATTGAGCGCAGGCCCGAC |
| CTH-ChIP-Region1 | GACGTTTCAGGCAACGCCTC | AGTGAAGCTGGAGTCCCTGG |

**Table S9. Primers used for real-time PCR for CTH and actin**

| Primer name | Forward Primer (5′ to 3′) | Reverse Primer (5′ to 3′) |
|---|---|---|
| RT-hCTH | GATTCGAAAGCCTTGCTGAG | ACTTCCACTTGGAGGGTGTG |
| RT-hActin | CTGGTGCCTGGGGCG | AGCCTCGCCTTTGCCGA |

**Table S10. Primers used for miRNA cDNA synthesis and real-time PCR experiments**

| Primer name | Sequence (5′ to 3′) | |
|---|---|---|
| SL-miR-150-5p | GTCGTATCCAGTGCAGGGTCCGAGGTATCGCACTGGATACGACCACTGGT | |
| SL-miR-510-5p | GTCGTATCCAGTCCAGGGTCCGAGGTATCGCACTGGATACGACTGATTG | |
| SL-U6 | GTCGTATCCAGTGCAGGGTCCGAGGTATTCGCACTGGATACGACTATGGAAC | |
| **Primer name** | **Forward Primer (5′ to 3′)** | **Reverse Primer (5′ to 3′)** |
| RT-miR-150-5p | GCGGCGGTCTCCCAACCCTTGT | GTGCAGGGTCCGAGGT |
| RT-miR-510-5p | GCGGCGGTACTCAGGFGAGTGG | GTGCAGGGTCCGAGGT |
| RT-U6 | CTGCGCAAGGATGACACGCA | GTGCAGGGTCCGAGGT |

**Table S11. mRNA levels (TPM) of CTH in pre- and post-transplanted human liver tissues-collected from Geodata set (accession: GSE151648)***

| S. no. | Sample ID | Pre-transplant | Post-transplant |
|---|---|---|---|
| 1 | GSM4587525 | 34.17 | 24.01 |
| 2 | GSM4587527 | 42.89 | 112.5 |
| 3 | GSM4587529 | 28.12 | 50.95 |
| 4 | GSM4587531 | 19.85 | 34.4 |
| 5 | GSM4587533 | 20.45 | 10.53 |
| 6 | GSM4587535 | 22.73 | 41.39 |
| 7 | GSM4587537 | 33.34 | 62.43 |
| 8 | GSM4587539 | 59.32 | 96.66 |
| 9 | GSM4587541 | 16.56 | 60.88 |
| 10 | GSM4587543 | 23.6 | 49.74 |
| 11 | GSM4587545 | 16.81 | 24.9 |
| 12 | GSM4587547 | 61.25 | 32.88 |
| 13 | GSM4587549 | 40.21 | 57.88 |
| 14 | GSM4587551 | 39.44 | 111 |
| 15 | GSM4587553 | 34.01 | 50.18 |
| 16 | GSM4587555 | 118.8 | 153.7 |
| 17 | GSM4587557 | 20.18 | 17.78 |
| 18 | GSM4587559 | 19.62 | 37.66 |
| 19 | GSM4587561 | 14.19 | 18.83 |
| 20 | GSM4587563 | 20.33 | 28.35 |
| 21 | GSM4587565 | 49.85 | 96.18 |
| 22 | GSM4587567 | 20.67 | 30.29 |
| 23 | GSM4587569 | 31.44 | 54.32 |
| 24 | GSM4587571 | 50.27 | 38.08 |
| 25 | GSM4587573 | 36.35 | 145.6 |
| 26 | GSM4587575 | 50.17 | 77.12 |
| 27 | GSM4587577 | 27.28 | 17.61 |
| 28 | GSM4587579 | 21.92 | 39.19 |
| 29 | GSM4587581 | 32.93 | 56.36 |
| 30 | GSM4587583 | 22.38 | 41.99 |
| 31 | GSM4587585 | 44.72 | 51.06 |
| 32 | GSM4587587 | 37.48 | 74.8 |
| 33 | GSM4587589 | 29.08 | 59.81 |
| 34 | GSM4587591 | 20.46 | 19.81 |
| 35 | GSM4587593 | 5.114 | 39.29 |
| 36 | GSM4587595 | 51.76 | 57.73 |
| 37 | GSM4587597 | 27.85 | 57.82 |
| 38 | GSM4587599 | 5.896 | 35.38 |
| 39 | GSM4587601 | 39.12 | 32.92 |
| 40 | GSM4587603 | 31.83 | 17.26 |

* The data was obtained from the NIH portal (https://www.ncbi.nlm.nih.gov/geo/query/acc.cgi?acc=GSE151648). The table shows transcript per million (TPM) values for CTH gene in pre- and post-transplanted human liver tissues.

**Table S12. mRNA levels (TPM) of CTH, CREB, AP-2α, NFY-A, and Sp1 in various human tissues-collected from GTex portal***

| Tissues | CTH | AP-2α | Sp1 | CREB | NFY-A |
|---|---|---|---|---|---|
| Adipose - Subcutaneous | 3.6825 | 0.21615 | 40.84 | 13.95 | 17.535 |
| Adipose - Visceral (Omentum) | 4.287 | 0.1904 | 31.62 | 11.31 | 16.98 |
| Adrenal Gland | 2.788 | 0.18695 | 24.875 | 5.7765 | 12.225 |
| Artery - Aorta | 4.163 | 0.6625 | 32.63 | 13.53 | 13.45 |
| Artery - Coronary | 3.968 | 0.3214 | 33.2 | 12.52 | 13.6 |
| Artery - Tibial | 5.518 | 0.663 | 39.06 | 14.84 | 13.33 |
| Bladder | 3.773 | 1.358 | 51.82 | 12.99 | 17.91 |
| Brain - Amygdala | 3.0025 | 0.09406 | 6.088 | 4.2385 | 9.055 |
| Brain - Anterior cingulate cortex (BA24) | 3.867 | 0.1221 | 5.179 | 4.468 | 8.223 |
| Brain - Caudate (basal ganglia) | 2.656 | 0.123 | 7.1555 | 4.125 | 9.1985 |
| Brain - Cerebellar Hemisphere | 4.232 | 2.5125 | 30.125 | 14.22 | 19.19 |
| Brain - Cerebellum | 4.733 | 2.632 | 30.76 | 10.31 | 18.48 |
| Brain - Cortex | 3.5275 | 0.13175 | 6.038 | 4.401 | 9.6485 |
| Brain - Frontal Cortex (BA9) | 4.484 | 0.1125 | 6.062 | 5.699 | 10.19 |
| Brain - Hippocampus | 2.346 | 0.145 | 6.265 | 4.589 | 9.963 |
| Brain - Hypothalamus | 4.121 | 0.0952 | 7.109 | 4.978 | 9.275 |
| Brain - Nucleus accumbens (basal ganglia) | 3.183 | 0.1124 | 6.831 | 4.593 | 9.621 |
| Brain - Putamen (basal ganglia) | 2.248 | 0.11705 | 5.809 | 3.5215 | 8.7125 |
| Brain - Spinal cord (cervical c-1) | 2.78 | 0.1944 | 13.56 | 7.872 | 17.47 |
| Brain - Substantia nigra | 3.167 | 0.064885 | 7.9185 | 4.216 | 10.53 |
| Breast - Mammary Tissue | 3.7925 | 14.855 | 38.075 | 12.97 | 18.855 |
| Cells - EBV-transformed lymphocytes | 18.08 | 0.1991 | 41 | 20.245 | 34.575 |
| Cells - Transformed fibroblasts | 7.073 | 0.3041 | 36.24 | 14.6 | 22.78 |
| Cervix - Ectocervix | 4.9605 | 1.08985 | 43.85 | 14.905 | 22.785 |
| Cervix - Endocervix | 4.619 | 1.485 | 52.12 | 17.22 | 25.71 |
| Colon - Sigmoid | 2.198 | 1.003 | 35 | 10.56 | 10.67 |
| Colon - Transverse | 3.8335 | 0.467 | 31.625 | 8.2005 | 11.605 |
| Esophagus - Gastroesophageal Junction | 2.871 | 0.89235 | 31.565 | 9.648 | 11.895 |
| Esophagus - Mucosa | 2.575 | 26.05 | 48.59 | 9.005 | 14.42 |
| Esophagus - Muscularis | 2.389 | 0.7789 | 30.995 | 9.2255 | 10.575 |
| Fallopian Tube | 7.574 | 0.4299 | 53.43 | 13.47 | 20.79 |
| Heart - Atrial Appendage | 1.367 | 0.4061 | 10.88 | 4.546 | 7.715 |

| | | | | | |
|---|---|---|---|---|---|
| Heart - Left Ventricle | 0.7716 | 0.08219 | 7.676 | 3.333 | 4.442 |
| Kidney - Cortex | 4.324 | 5.066 | 19.61 | 4.228 | 7.638 |
| Liver | 44.92 | 0.0537 | 10.51 | 2.821 | 7.249 |
| Lung | 4.403 | 0.2445 | 37.83 | 12.9 | 20.91 |
| Minor Salivary Gland | 2.951 | 20.59 | 38.78 | 9.985 | 13.63 |
| Muscle - Skeletal | 1.6245 | 0.042155 | 17.98 | 3.0515 | 6.676 |
| Nerve - Tibial | 8.4145 | 0.9261 | 48.305 | 14.62 | 20.97 |
| Ovary | 21.03 | 0.2239 | 43.27 | 14.28 | 31.43 |
| Pancreas | 5.063 | 0.076215 | 12.315 | 3.6145 | 6.9575 |
| Pituitary | 3.989 | 0.3907 | 18.82 | 8.142 | 16.84 |
| Prostate | 4.673 | 10.42 | 31.295 | 10.925 | 16.955 |
| Skin - Not Sun Exposed (Suprapubic) | 1.414 | 61.58 | 44.58 | 10.65 | 14.16 |
| Skin - Sun Exposed (Lower leg) | 1.61 | 61.2 | 46.6 | 10.72 | 13.62 |
| Small Intestine - Terminal Ileum | 2.495 | 0.274 | 34.08 | 9.912 | 13.71 |
| Spleen | 1.5395 | 0.14125 | 35.695 | 12.955 | 19.305 |
| Stomach | 6.2025 | 0.327 | 21.665 | 6.007 | 11.21 |
| Testis | 8.578 | 2.047 | 34.81 | 16.81 | 62.8 |
| Thyroid | 10.44 | 0.6096 | 35.83 | 13.075 | 29.5 |
| Uterus | 5.168 | 0.2544 | 56.9 | 15.37 | 23.58 |
| Vagina | 2.693 | 20 | 43.2 | 13.25 | 17.01 |
| Whole Blood | 0.3549 | 0.01534 | 39.65 | 5.833 | 13.59 |

* The data was obtained from GTEx portal (https://www.gtexportal.org/home/). The table shows transcript per million (TPM) values for CTH, AP-2α, Sp1, CREB and NFY-A genes from various tissues.

**Table S13. mRNA levels (TPM) of CTH, CREB, AP-2α, NFY-A, and Sp1 in human liver tissue-collected from Geodata set (accession: GSE151648)***

| S no. | Sample | CTH | CREB | AP-2α | NFY-A | Sp1 |
|---|---|---|---|---|---|---|
| 1 | GSM4587525 | 34.17 | 1.77 | 0.07966 | 3.221 | 9.942 |
| 2 | GSM4587527 | 42.89 | 1.839 | 0 | 2.647 | 11.04 |
| 3 | GSM4587529 | 28.12 | 2.192 | 0.05237 | 4.765 | 12.93 |
| 4 | GSM4587531 | 19.85 | 1.572 | 0.07042 | 2.855 | 9.495 |
| 5 | GSM4587533 | 20.45 | 2.372 | 0.1733 | 3.091 | 11.65 |
| 6 | GSM4587535 | 22.73 | 1.826 | 0 | 3.002 | 10.73 |
| 7 | GSM4587537 | 33.34 | 2.136 | 0.02734 | 3.536 | 9.964 |
| 8 | GSM4587539 | 59.32 | 1.504 | 0 | 3.179 | 11.16 |
| 9 | GSM4587541 | 16.56 | 1.767 | 0.1031 | 2.825 | 10.09 |
| 10 | GSM4587543 | 23.6 | 1.592 | 0.02638 | 2.82 | 9.612 |
| 11 | GSM4587545 | 16.81 | 1.19 | 0.0816 | 2.456 | 5.151 |
| 12 | GSM4587547 | 61.25 | 2.303 | 0.0282 | 4.5 | 14.18 |
| 13 | GSM4587549 | 40.21 | 1.848 | 0.08212 | 3.292 | 9.313 |
| 14 | GSM4587551 | 39.44 | 1.744 | 0 | 3.253 | 12.56 |
| 15 | GSM4587553 | 34.01 | 2.063 | 0.1216 | 3.322 | 9.658 |
| 16 | GSM4587555 | 118.8 | 1.892 | 0 | 2.759 | 10.13 |
| 17 | GSM4587557 | 20.18 | 1.545 | 0 | 2.799 | 9.858 |
| 18 | GSM4587559 | 19.62 | 1.922 | 0.0119 | 3.602 | 9.298 |
| 19 | GSM4587561 | 14.19 | 1.27 | 0.05196 | 2.091 | 6.812 |
| 20 | GSM4587563 | 20.33 | 1.608 | 0.01189 | 4.569 | 8.776 |
| 21 | GSM4587565 | 49.85 | 1.572 | 0.03347 | 2.938 | 8.493 |
| 22 | GSM4587567 | 20.67 | 1.96 | 0.0916 | 3.249 | 11.41 |
| 23 | GSM4587569 | 31.44 | 2.013 | 0.07244 | 2.99 | 10.4 |
| 24 | GSM4587571 | 50.27 | 2.282 | 0 | 4.479 | 12.9 |
| 25 | GSM4587573 | 36.35 | 1.865 | 0.02741 | 3.717 | 10.6 |
| 26 | GSM4587575 | 50.17 | 2.075 | 0.0901 | 3.984 | 11.06 |
| 27 | GSM4587577 | 27.28 | 2.059 | 0.01999 | 3.395 | 11.26 |
| 28 | GSM4587579 | 21.92 | 2.063 | 0.1649 | 4.069 | 11.6 |
| 29 | GSM4587581 | 32.93 | 1.268 | 0.09943 | 2.527 | 8.493 |
| 30 | GSM4587583 | 22.38 | 2.312 | 0.1113 | 5.008 | 13.88 |
| 31 | GSM4587585 | 44.72 | 2.675 | 0.04795 | 4.363 | 14.1 |
| 32 | GSM4587587 | 37.48 | 1.762 | 0.02648 | 3.425 | 9.066 |
| 33 | GSM4587589 | 29.08 | 1.96 | 0.1188 | 3.48 | 13.23 |
| 34 | GSM4587591 | 20.46 | 2.08 | 0.03983 | 3.1 | 9.711 |
| 35 | GSM4587593 | 5.114 | 0.4731 | 0 | 0.7419 | 1.376 |

| | | | | | | |
|---|---|---|---|---|---|---|
| 36 | GSM4587595 | 51.76 | 1.594 | 0.05763 | 3.5 | 10.59 |
| 37 | GSM4587597 | 27.85 | 2.11 | 0.02795 | 3.636 | 12.89 |
| 38 | GSM4587599 | 5.896 | 0.7416 | 0.04251 | 0.7353 | 2.32 |
| 39 | GSM4587601 | 39.12 | 2.029 | 0 | 4.268 | 10.92 |
| 40 | GSM4587603 | 31.83 | 1.415 | 0.05487 | 3.516 | 7.137 |

* The data was obtained from the NIH portal (https://www.ncbi.nlm.nih.gov/geo/query/acc.cgi?acc=GSE151648). The table shows transcript per million (TPM) values for CTH, AP-2α, Sp1, CREB and NFY-A genes in liver tissues.

### Supplementary Figures

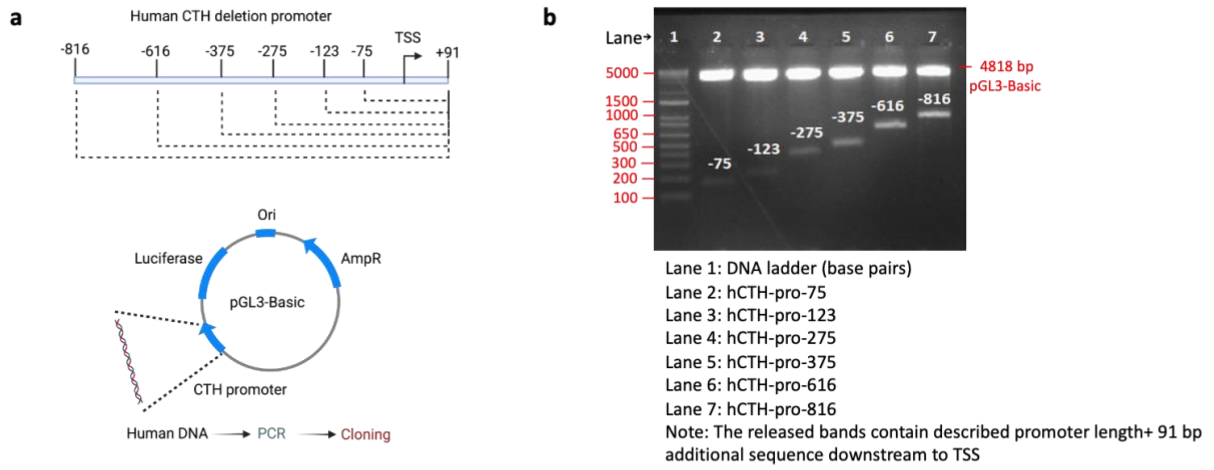


**Figure S1. Schematic representation of promoter cloning and promoter-reporter clone confirmation.** (a) Varying lengths of human CTH promoters were PCR amplified and ligated to the pGL3-Basic promoter less luciferase reporter vector. (b) The clones were confirmed by checking the release of deleted promoter inserts on agarose gel following restriction digestion using *BamHI* and *Kpn I* restriction enzymes. The sequences of promoter inserts were confirmed by Sanger DNA sequencing.

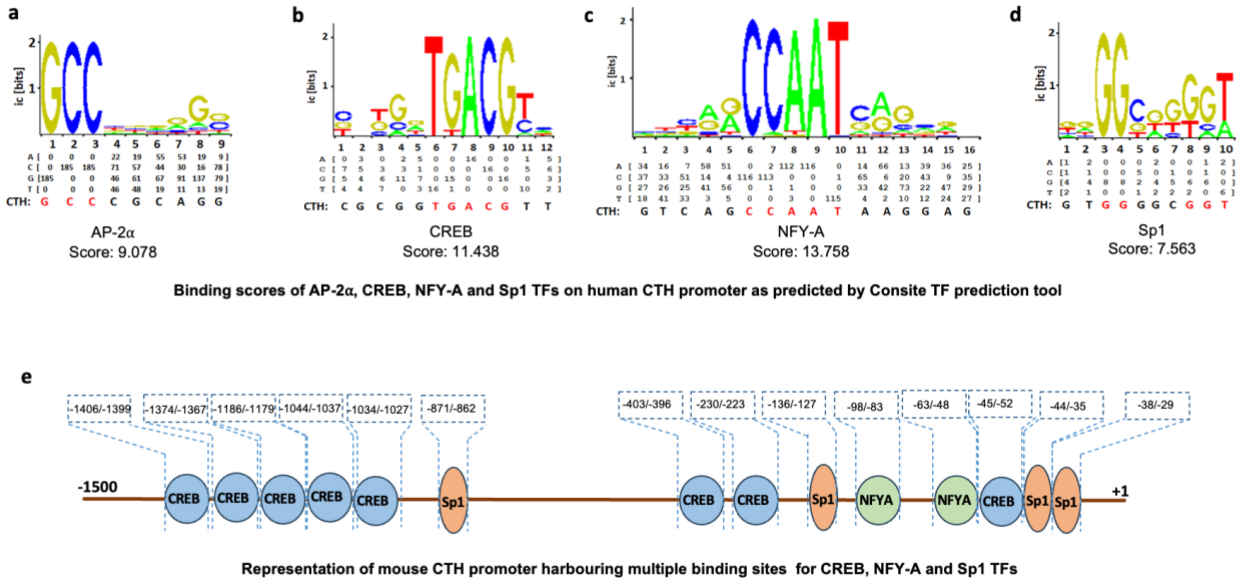


**Figure S2. TF binding scores for human CTH promoter and TF binding site conservation in mouse CTH promoter.** (a-d) The binding scores were predicted by ConSite which is based on the similarity between the predicted binding site and position weight matrix of AP-2α, CRE, Sp1, and NFY-A. (e) Of all the four TFs, the binding sites for CREB, Sp1, and NFY-A (but not for AP-2α) were found in the mouse CTH promoter. Multiple binding sites were predicted for CREB (8 sites), Sp1 (4 sites), and NFY-A (2 sites) within -1500 bp promoter region of mouse CTH gene.

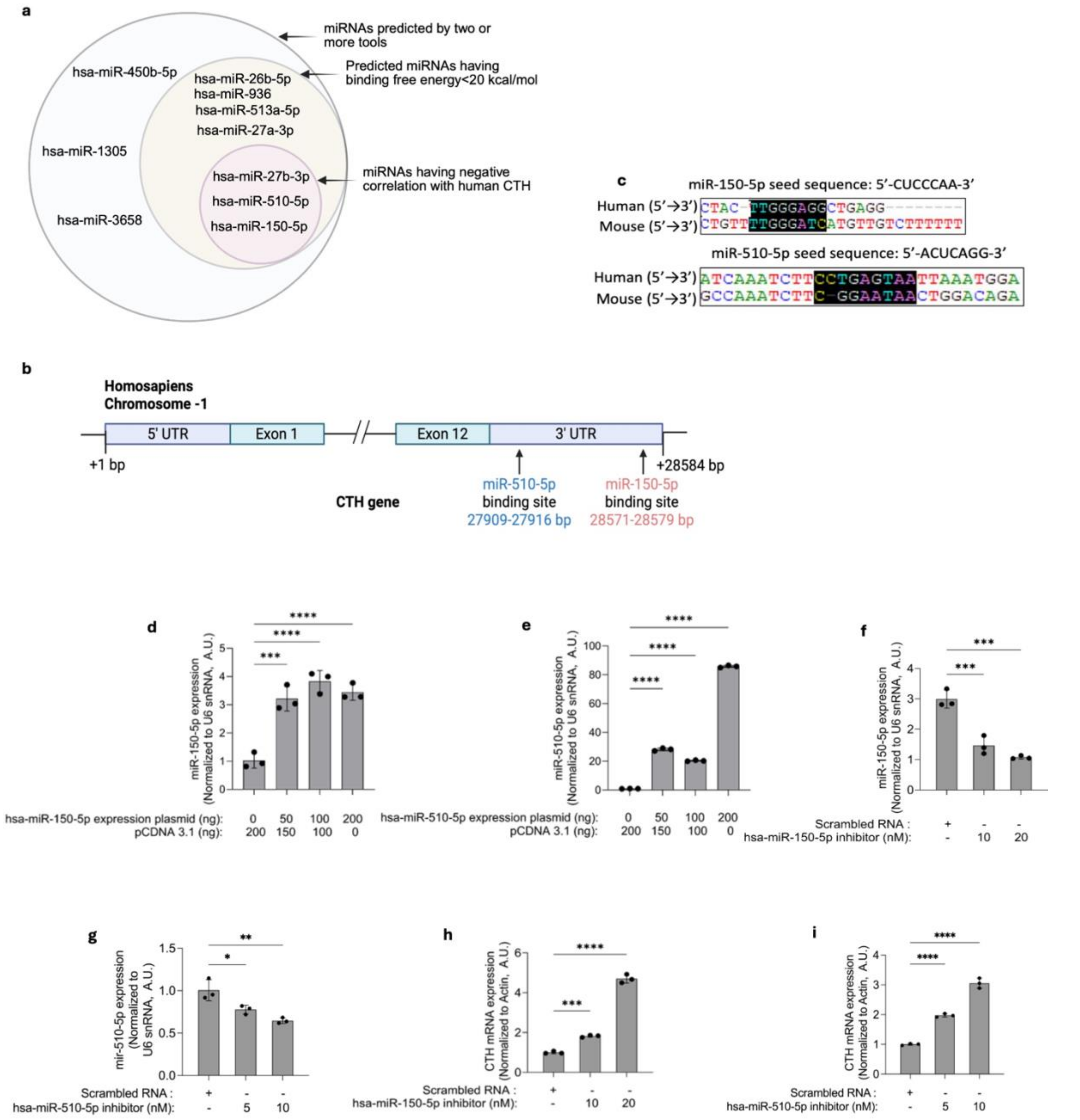


**Figure S3. Regulatory role of miR-150-5p and miR-510-5p on CTH expression.** (a) Venn diagram showing different miRNAs that were shortlisted using parameters such as redundancy of prediction by miRNA prediction tools, followed by binding free energy and their expression correlation with CTH. (b) Schematic representation of the predicted binding sites of miR-150-5p and miR-510-5p in 3'-UTR of human CTH. (c) Binding sites for miR-150-5p and miR-510-5p is conserved in mouse CTH 3'-UTR (c). (d and e) Dose-dependent increase in the expression of miR-150-5p and miR-510-5p with increasing doses of hsa-miR-150-5p and hsa-miR-510-5p expression plasmids in Huh7 cells, respectively. (f-i) Decrease in the endogenous miR-150-5p and miR-510-5p levels followed by increase in CTH mRNA levels were observed after treatment of Huh7 cells with hsa-miR-150-5p and hsa-miR-510-5p inhibitors, respectively. The statistical significance was determined by one-way ANOVA with Tukey's multiple correction as a post-hoc test. ****P<0.0001, ***P<0.001, **P<0.01, *P<0.05.

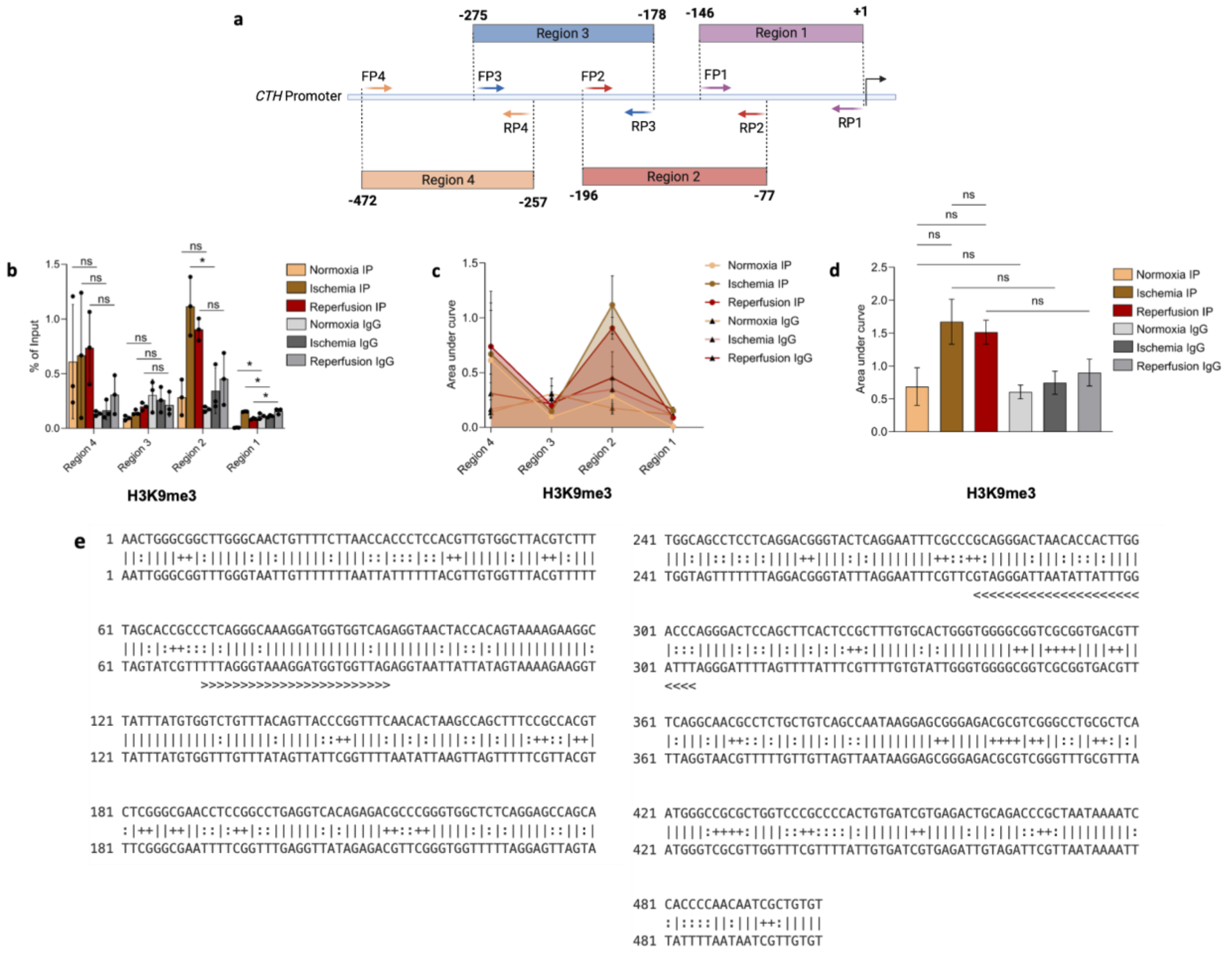


**Figure S4. Chromatin modifications in the CTH promoter region.** (a) ChIP assays using antibody against H3K9me3 followed by qPCR for the enrichment of CTH promoter was performed to understand the role of epigenetic modification in CTH expression during normoxia, ischemia and reperfusion conditions. Schematic diagram representing the strategy to quantify the enriched promoter after ChIP. (b) Bar plot showing the enrichment of different regions of human CTH promoter associated with H3K9me3 under normoxia, ischemia, and reperfusion conditions. (c) Line plot showing the area under curve calculated from bar plot panel b for each region of the promoter. (d) Cumulative area of all the regions of CTH promoter plotted as bar plot for normoxia, ischemia and reperfusion. (e) CpG sites (total 32 sites) in 500 bp human CTH promoter as predicted by MethPrimer tool. The statistical significance was determined by one-way ANOVA with Tukey's multiple correction as a post-hoc test. *P<0.05; ns: not significant.
